# Pre-mRNA splicing inhibits m^6^A deposition, allowing longer mRNA half-life and flexible protein coding

**DOI:** 10.1101/2022.12.26.521933

**Authors:** Zhiyuan Luo, Qilian Ma, Shan Sun, Ningning Li, Hongfeng Wang, Zheng Ying, Shengdong Ke

## Abstract

Both pre-mRNA splicing and *N*^6^-methyladenosine (m^6^A) mRNA modification occur during transcription, enabling the potential crosstalk regulation between these two fundamental processes. The regional m^6^A location bias of avoiding splice site region, calls for an open hypothesis whether pre-mRNA splicing could affect m^6^A deposition. By deep learning modeling, we find that pre-mRNA splicing represses a proportion (4% to 32%) of m^6^A deposition at nearby exons. Experimental validation confirms such an inhibition as the m^6^A signal increases in mRNA once the host gene does not undergo pre-mRNA splicing to produce the same mRNA. Pre-mRNA splicing inhibited m^6^A sites tend to have higher m^6^A enhancers and lower m^6^A silencers locally than the m^6^A sites that are not inhibited. Moreover, this m^6^A deposition inhibition by pre-mRNA splicing shows high heterogeneity at different exons of mRNAs at genome-widely, with only a small proportion (12% to 15%) of exons showing strong inhibition, enabling stable mRNAs and flexible protein coding for important biological functions.

## Introduction

As the most abundant mRNA internal modification, the *N*^6^-methyladenosine (m^6^A) is involved in various biological processes including cell differentiation, brain development, tumorigenesis^1–6^, and could affect multiple aspects of RNA metabolism, including transcription, splicing, translation, and degradation^7,8^, with a major function in promoting mRNA decay^9–12^. The m^6^A is deposited to nascent pre-mRNA co-transcriptionally^11^, primarily by the <u>m</u>ethyl<u>t</u>ransferase <u>c</u>omplex (MTC) comprising the catalytic core METTL3-METTL14 heterodimer and other factors^13–19^. m^6^A is installed at a motif consensus of RR<u>A</u>CH (R = A or G, H = A, C, or U) as a stringent motif or R<u>A</u>C as a more inclusive motif^20–22^. Despite the wide prevalence of m^6^A consensus in mRNA, only a very small fraction is methylated^11,20^. At the global level, m^6^As reside preferentially in last exons, as well as in long internal exons^11,20^. Furthermore, m^6^As in internal exons appear to avoid the nearby exonic region close to splice sites^11^. Our previous work has revealed that the m^6^A site-specific methylation was primarily determined by the flanking nucleotide sequences, and the local functional *cis*-elements mainly resided within the 50 nt downstream of the site^23^. The underlying mechanism beyond the identification of local *cis*-regulatory elements of m^6^A site-specificity is still largely unknown.

As with m^6^A deposition, pre-mRNA splicing is also coupled with transcriptional events, allowing for potential functional crosstalk during transcription. Though several studies suggested that m^6^A could regulate alternative splicing^21,24–27^, a careful bioinformatics analysis showed that loss of METTL3 in mouse embryonic stem cells had a minimal effect on pre-mRNA splicing^11^. Conversely, whether pre-mRNA splicing could affect m^6^A deposition is an open question. Most m^6^A deposition occur in the region moving away from last exon start and appears to avoid the adjacent region close to splice sites in internal exons^11,20^. These m^6^A regional distribution biases suggest that pre-mRNA splicing could potentially play an inhibitory role for the m^6^A deposition at the nearby region close to splice sites.

Previously we have established the iM6A deep learning model which models m^6^A site specificity with high accuracy (AUROC=0.99) by using the primary nucleotide sequence flanking the m^6^A site^23^. This work demonstrated that the site specificity of m^6^A modification was encoded primarily by the flanking nucleotide sequence at the *cis*-level. Though the deep learning model itself is hard to be understood directly (i.e. a “black box”), we could probe for the underlying biological insights by creative in-silico mutation of natural genomic regions to test our hypotheses. Then if the followed wet experiments validate randomly selected simulations, this contributes to verifying the model and the biological hypotheses it is designed to investigate. As an initial study, we performed the *in silico* saturation mutagenesis on the local sequences surrounding the m^6^A site and discovered that the downstream 50 nt region of the m^6^A site was highly enriched with the *cis*-elements governing m^6^A deposition^23^. Independent experimental validation supported this finding. The *in silico* deep learning modeling approach has proved to be an effective way to investigate the *cis*-regulatory mechanisms that determines m^6^A deposition, and offers a high-throughput and fast-paced low-cost discovery mechanism relative to exclusively experimental studies which could be cost-prohibitive^23^.

In this study, we implemented iM6A deep learning modeling to investigate *cis*-regulatory mechanisms for m^6^A site specificity beyond the local *cis*-regulatory elements. By the *in silico* mutational modeling at intron deletion in genes, we discovered that pre-mRNA splicing inhibits a proportion of m^6^A deposition at nearby exons. These splicing-inhibited m^6^A sites tended to have a good local *cis*-element environment with more m^6^A enhancers and fewer m^6^A silencers, compared to the m^6^A sites that were not inhibited. These modeling findings were supported by the experimental validation, as will be shown in Fig. 3. The m^6^A deposition inhibition by pre-mRNA splicing exhibited a high heterogeneity at genomic level, with a small proportion of exons exhibiting strong inhibition. By this m^6^A deposition inhibition mechanism by pre-mRNA splicing, multi-exon mRNA will have longer half-life given the same primary nucleotide sequence; Also this mechanism allows mRNA to encode protein sequence flexibly with less concern of creating too many m^6^A sites to compromise its mRNA stability.

**Fig. 1:**
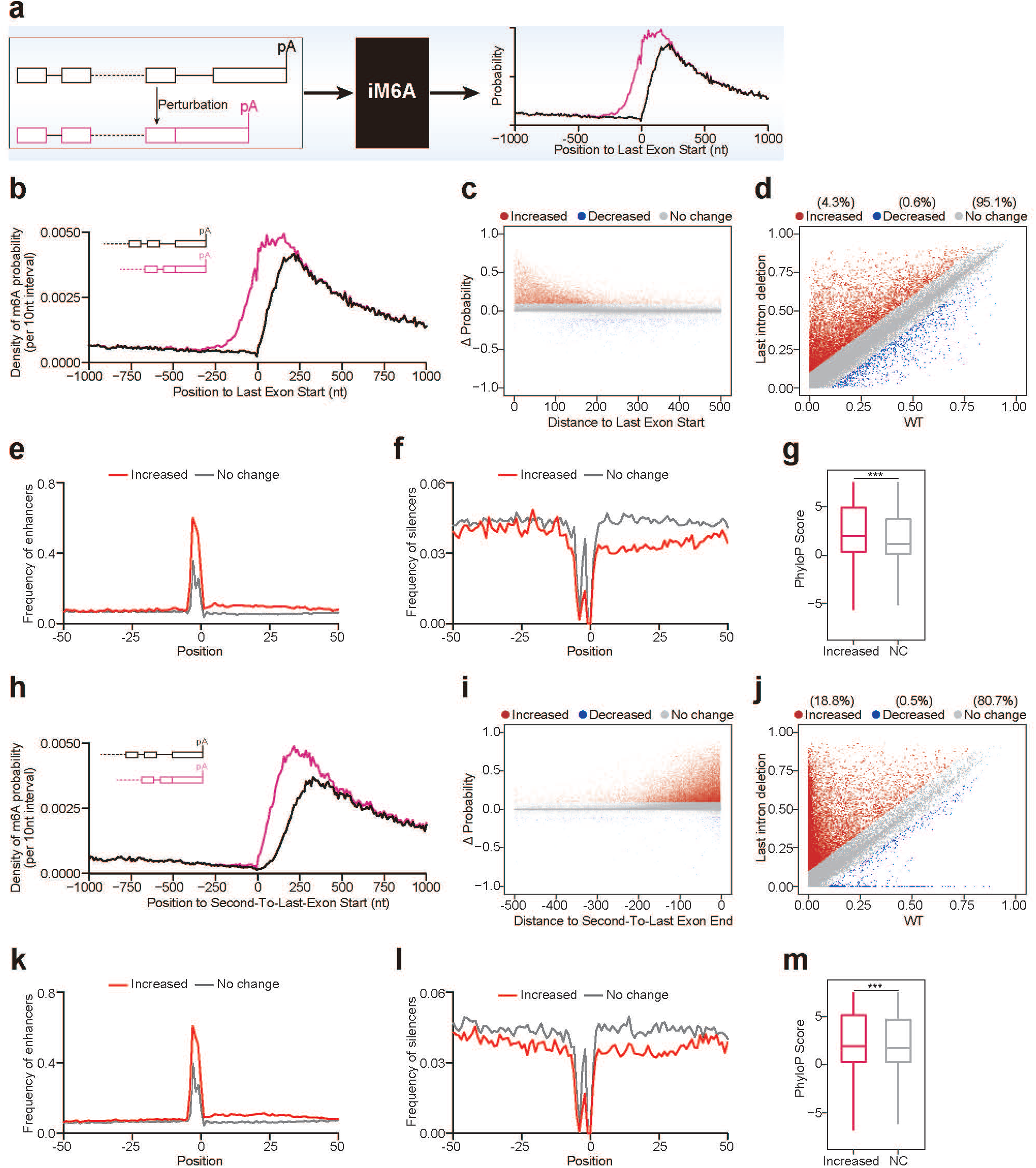
Deep learning modeling reveals last intron deletion revives m^6^A deposition at last exon and second-to-last exon. **a**, Schematic figure of the deep learning modeling m^6^A deposition in pre-mRNA by iM6A. The nucleotide sequence of pre-mRNA served as input, while the output of iM6A was the probability value of each nucleotide being a m^6^A site. **b**, The m^6^A density of transcripts around last exon start were compared between full length (black line) and last intron deletion (pink line). The value was calculated as the total probability value in a 10-nt interval divided by the total number of mRNAs in the interval. **c-d**, Positional plot of ΔProbability (Fig. 1c) and scatter plot of probability (Fig. 1d) for the RAC sites located in last exons. Red dots (Increased: n = 7033 for 4000 genes), blue dots (Decreased: n = 911 for 4000 genes), and grey dots (no change: n = 155682 for 4000 genes) were those sites that had increased probability (> 0.1), decreased probability (< - 0.1), or not change probability (|ΔProbability| <= 0.1) respectively by last intron deletion. **e-f**, Positional plot for the frequency of top 50 enhancers (Fig. 1e), silencers (Fig. 1f) in mRNA sequences around the RAC sites. The sites were located in last exons, and the plots were compared between the increased sites (red line, ΔProbability > 0.1) and no change sites (grey line, |ΔProbability| <= 0.1). **g**, Box plot of PhyloP score of latent m^6^A sites or no change sites in last exons (n = 19453 or 143022). Median and interquartile ranges were presented for the box plot. The p-values were calculated by Wilcoxon test (Significance: *** p < 0.001). **h**, The m^6^A density of transcripts around second-to-last exon start were compared between full length (black line) and last intron deletion (pink line). **i-j**, Positional plot of ΔProbability (Fig. 1i) and scatter plot of probability (Fig. 1j) for the RAC sites located in second-to-last exons. Red dots (Increased: n = 16563 for 16769 genes), blue dots (Decreased: n = 446 for 16769 genes), and grey dots (no change: n = 71323 for 16769 genes) were those sites that had increased probability (> 0.1), decreased probability (< −0.1), or not change probability (|ΔProbability| <= 0.1) respectively by last intron deletion. **k-l,** Positional plot for the frequency of top 50 enhancers (Fig. 1k), silencers (Fig. 1l) in mRNA sequences around the RAC sites. The sites were located in second-to-last exon, and the plots were compared between the increase sites (red line, ΔProbability > 0.1) and no change sites (grey line, |ΔProbability| <= 0.1). **m**, Box plot of PhyloP score of latent m^6^A sites or no change sites in second-to-last exons (n = 14039 or 54906). Median and interquartile ranges were presented for the box plot. The p-values were calculated by Wilcoxon test (Significance: *** p < 0.001).

**Fig. 2:**
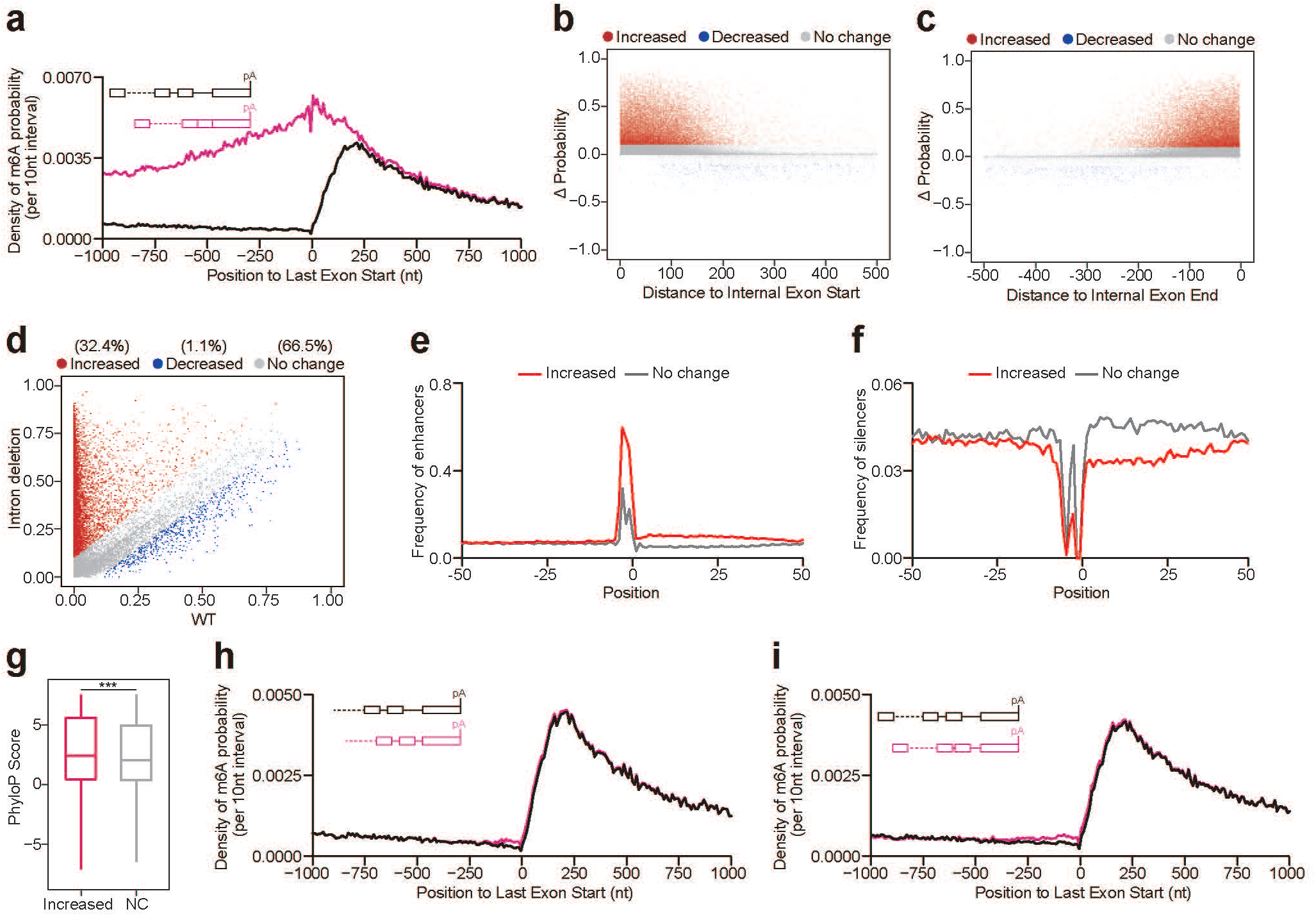
Deep learning modeling reveals introns deletion revives m^6^A deposition at internal exons. **a**, The m^6^A density of transcripts around last exon start were compared between full length (black line) and all introns deletion (pink line). **b-d**, Positional plot of ΔProbability (Fig. 1b,c) and scatter plot of probability (Fig. 1d) for the RAC sites located in internal exons. Red dots (Increased: n = 20730 for 1500 genes), blue dots (Decreased: n = 682 for 1500 genes), grey dots (no change: n = 42509 for 1500 genes) were those sites that had increased probability (> 0.1), decreased probability (< − 0.1), or not change probability (|ΔProbability| <= 0.1) respectively by all introns deletion. **e-f**, Positional plot for the frequency of top 50 enhancers (Fig. 1e), silencers (Fig. 1f) in mRNA sequences around the RAC sites. The sites were located in internal exons, and the plots were compared between the increase sites (red line, ΔProbability > 0.1) and no change sites (grey line, |ΔProbability| <= 0.1). **g**, Box plot of PhyloP score of latent m^6^A sites or no change sites in internal exons (n = 209292 or 429473). Median and interquartile ranges were presented for the box plot. The p-values were calculated by Wilcoxon test (Significance: *** p < 0.001). **h-i**, The m^6^A density of transcripts around last exon start were compared between full length (black line) and intron truncation (pink line) (Fig. h for last intron truncation, Fig. 1i for all introns truncation). The sequences of introns (> 400 nt) were truncated to 400 nucleotides by keeping 200 nucleotides of intron start and intron end.

**Fig. 3:**
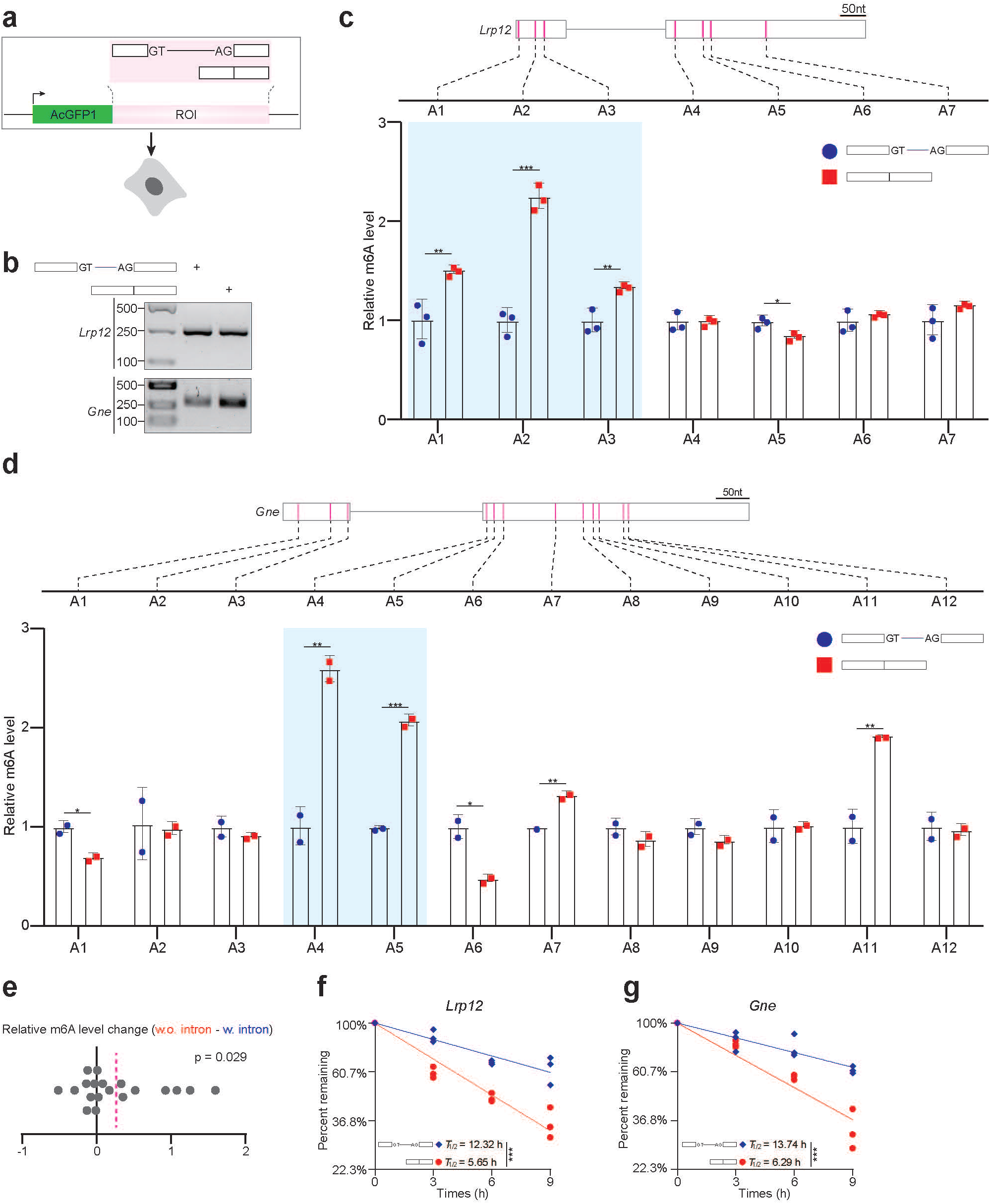
Experimental validation of intron repression on m^6^A deposition. **a**, Illustration of the minigene experiment, using *Lrp12* and *Gne* as two model genes. The minigene contained two exons (100 nt long upstream exon and 240_nt long downstream exon, details in Methods) and a 200 nt intervening mini-intron in between, and the first exon was in-frame fused to AcGFP1. Constructs were transfected into HEK-293T cells, and m^6^A signal was quantified by SELECT method. **b**, The pre-mRNA splicing of mini-intron in minigene of *Lrp12* or *Gne* was validated by RT-PCR in HEK-293T cells. **c-d**, The bar plot of relative m^6^A level for detecting the m^6^A sites in mRNA. The constructs of minigenes were shown, and RAC sites in *Lrp12* (Fig. 3c) or *Gne* (Fig. 3d) were marked as pink lines. Data were presented as mean ± SD, the p-values were calculated by Student t-test (Significance: *** p < 0.001, ** p < 0.01, * p < 0.05). The RAC site showed agreed increased m^6^A signal in both iM6A modeling and experimental validation was marked by blue box. **e**, The dot plot of the experimental determined relative m^6^A level change for each RAC site in mRNA (19 sites in total). Relative m^6^A level change was calculated for the relative m^6^A level of each site between intron-containing and intron-deletion mRNAs. The mean value (0.264) was shown as the dotted pink line, and p-value was calculated by one-sample t-test for the increase vs. no change. **f-g**, mRNA decay plotted as a function of time. The normalized levels of minigene mRNA (Fig. 3f for *Lrp12*, and Fig. 3g for *Gne*) at 0 h were set to 100%. The y axis represents the log value of mRNA remaining level. The p-values were calculated by Student t-test (Significance: *** p < 0.001).

## Results

### Deep learning modeling revealed that pre-mRNA splicing inhibits m^6^A deposition at last exon and second-to-last exon

As we previously found that m^6^A appeared to avoid the nearby region close to splice sites while being mostly enriched in the region moving away from last exon starts^11,20^, we speculated that pre-mRNA splicing might inhibit m^6^A deposition at exons. We modeled this with an *in silico* mutational experiment by deleting the last intron sequences from the gene to generate the non-last intron genes as the input for iM6A (Fig. 1a) (i.e. pre-mRNA would not undergo pre-mRNA splicing of last intron to generate mRNA). We unexpectedly found that the m^6^A density increased around last exon start (Fig. 1b for mouse, and Supplementary Fig. 1a for human). A more detailed examination down to individual RAC sites in this region revealed that 1) a proportion of RAC sites (~4%) in last exons had an increase in m^6^A deposition (Fig. 1d for mouse, and Supplementary Fig. 1c for human). Since the m^6^A deposition of these sites were repressed by the pre-mRNA splicing of last intron, we define them as the splicing-repressed m^6^A sites or latent m^6^A sites; 2) most of those sites were enriched within the ~100 nt region to last exon start (Fig. 1c for mouse, and Supplementary Fig. 1b for human). Given that the local *cis*-elements that govern m^6^A deposition largely reside within the 50 nt downstream of the sites where we previously identified m^6^A enhancers and m^6^A silencers^23^, we further investigated the distribution of m^6^A enhancers and m^6^A silencers in the local region flanking the RAC sites upon last intron deletion. In comparison to the majority RAC sites without m^6^A deposition change, the RAC sites with increased m^6^A deposition contained more m^6^A enhancers in the downstream 50 nt region (Fig. 1e for mouse, and Supplementary Fig. 1d for human) while hosting less m^6^A silencers in the same region (Fig. 1f for mouse, and Supplementary Fig. 1e for human). This data showed that those latent m^6^A sites (ΔProbability > 0.1) in last exons had a favorable local *cis*-element composition for m^6^A deposition but was repressed by pre-mRNA splicing. Evolution conservation analysis showed that these splicing-repressed m^6^A sites were more conserved in comparison to the RAC sites that were not subject to this pre-mRNA splicing inhibition (Fig. 1g for mouse, and Supplementary Fig. 1f for human), supporting their functional importance.

Besides repressing the m^6^A deposition in last exons, pre-mRNA splicing might also inhibit the m^6^A deposition in the second-to-last exons (Fig. 1h for mouse, and Supplementary Fig. 1g for human). To comprehensively investigate this hypothesis, we plotted the detailed m^6^A methylation changes for all the RAC sites in the second-to-last exons. Upon the last intron deletion, ~19% RAC sites had increased m^6^A probability (Fig. 1j for mouse, and Supplementary Fig. 1i for human), and most of those latent sites were also enriched in the ~100 nt region close to the end of second-to-last exons (Fig. 1i for mouse, and Supplementary Fig. 1h for human). Similarly, the m^6^A enhancers enriched and m^6^A silencers avoided in the 50 nt downstream region of these latent m^6^A sites respectively (Fig. 1k,l for mouse, and Supplementary Fig. 1j,k for human). These data demonstrated that pre-mRNA splicing inhibits the local m^6^A deposition at its two adjacent exons while not affecting other upstream exons (Fig. 1h for mouse, and Supplementary Fig. 1g for human). In addition, these splicing-repressed m^6^A sites were also more conserved in comparison to the RAC sites that were not subject to this intron inhibition suggesting their functional importance (Fig. 1m for mouse, and Supplementary Fig. 1l for human).

### Deep learning modeling revealed that pre-mRNA splicing inhibits m^6^A deposition at internal exons

Since the last and internal exons have the same *cis*-element code in regulating m^6^A deposition^23^, it is possible that pre-mRNA splicing also inhibits m^6^A deposition in internal exon. To validate this, we performed a new round of m^6^A deposition *in silico* modeling by deleting all introns from the gene (i.e. pre-mRNA would not undergo pre-mRNA splicing to generate mRNA), and found that the m^6^A level at internal exons also increased remarkably upon intron deletion (Fig. 2a,b,c for mouse, and Supplementary Fig. 2a,b,c for human). Overall ~32% RAC sites in internal exons showed higher m^6^A probability (Fig. 2d for mouse, and Supplementary Fig. 2d for human), and those latent m^6^A sites also mostly resided in the ~100 nt region to the two ends of internal exons (Fig. 2b,c for mouse, and Supplementary Fig. 2b,c for human). In addition, the m^6^A enhancers or silencers were enriched or avoided in the 50 nt downstream region of these splicing-repressed m^6^A sites respectively, again supporting that these splicing-repressed m^6^A sites had a good local *cis*-elements composition for m^6^A deposition but were repressed by the nearby pre-mRNA splicing (Fig. 2e,f for mouse, and Supplementary Fig. 2e,f for human). Evolution conservation analysis demonstrated that these splicing-repressed m^6^A sites were more conserved in comparison to the RAC sites that were not subject to this pre-mRNA splicing inhibition (Fig. 2g for mouse, and Supplementary Fig. 2g for human).

To further understand the m^6^A inhibition by pre-mRNA splicing, we truncated either last intron (Fig. 2h for mouse, and Supplementary Fig. 2h for human) or all introns (Fig. 2i for mouse, and Supplementary Fig. 2i for human) to a maximum of 400 nucleotides by keeping the nearest 200 nucleotides at the two intron ends. As intronic splicing *cis*-elements are highly enriched at the 100 nt flanking intronic region of most human and mouse exons^28^, these mini-introns should mostly retain their splicing capacity. Intron size reduction only altered the m^6^A density mildly (Fig. 2h,i for mouse, and Supplementary Fig. 2h,i for human), suggesting that the deep intronic sequences only played a minor role in inhibiting m^6^A deposition at nearby exons. We further truncated the full-length last introns to 200 nucleotides mini-introns by preserving the flanking 100 nucleotides of the two intron ends which contain highly enriched intronic splicing *cis*-elements^28^ (Supplementary Fig. 3a,b,c). <u>A</u>s above, the deep intronic sequence contributed little to this m^6^A deposition inhibition (Fig. 2h,i, and Supplementary Fig. 2h,i), and the m^6^A density at the ends of the two flanking exons had little change upon this intron length truncation (Supplementary Fig. 3a,b,c). In contrast, the deletion of mini-introns promoted m^6^A deposition at ~100 nt region of the two nearby exons (Supplementary Fig. 3a,b,c). These data support that the pre-mRNA splicing of the 200 nt long mini-intron may be as potent in inhibiting m^6^A deposition at nearby exons as the pre-mRNA splicing of the full-length intron, enabling the minigene experimental validation below.

### Experimental validation of pre-mRNA splicing inhibition on m^6^A deposition

To experimentally validate the pre-mRNA splicing inhibition on m^6^A deposition, we ligated the <u>c</u>o<u>d</u>ing <u>s</u>equence (CDS) of AcGFP1 in-frame to a minigene. The minigene consisted of two exons and a 200 nt intervening mini-intron (Fig. 3a). We constructed two such minigenes, *Lrp12* and *Gne*. The pre-mRNA splicing of both minigenes occurred efficiently (Fig. 3b), experimentally confirming that the 200 nt long mini-intron retained its splicing capacity. The iM6A modeling predicted the m^6^A inhibition by pre-mRNA splicing in both minigenes, *Lrp12* and *Gne* (Supplementary Fig. 4a,b). Consistently, using the SELECT method to experimentally quantify m^6^A^29^, we did observe the m^6^A signal increase in both minigenes when they did not undergo pre-mRNA splicing to produce the mRNA with the same nucleotide sequence (Fig. 3c,d). Altogether, eight RAC sites were predicted to increase their m^6^A level when the minigene did not undergo pre-mRNA splicing to produce the mRNA with the same nucleotide sequence (predicted m^6^A level increase > 0.1) (Supplementary Fig. 4a), and five such RAC sites were experimentally confirmed to increase their m^6^A level (highlighted in Fig. 3c, d). We experientially quantified all 19 RAC sites both minigenes and found that they overall had an evident m^6^A signal increase (average relative m^6^A level increase = 0.264 > 0, p = 0.029, one sample t-test) (Fig. 3e), agreeing with the iM6A prediction (average predicted methylation level increase = 0.197 > 0, p = 0.0004, one sample t-test) (Supplemental Fig. 4b). These experimental data confirmed that pre-mRNA splicing inhibits m^6^A deposition at nearby exons (Fig. 3, and Supplementary Fig. 4). At the same time, we observed the RAC sites in individual nearby exons had distinct m^6^A deposition inhibition, some exons were strongly inhibited by pre-mRNA splicing, while others were not (Fig. 3c,d), suggesting heterogeneity of m^6^A deposition inhibition.

Since a major function of m^6^A is promoting mRNA decay^9–12^, the mRNA produced without pre-mRNA splicing inhibition has stronger m^6^A signal, and thus should have shorter half-life (*T*_1/2_). As expected, for both *Lrp12* and *Gne*, the mRNAs produced by constructs that didn’t undergo pre-mRNA splicing had shorter *T*_1/2_s than mRNAs produced by constructs that did undergo pre-mRNA splicing, though these two mRNAs shared identical primary nucleotide RNA sequence (Fig. 3f,g).

### A small proportion of last exons exhibit strong m^6^A deposition inhibition by pre-mRNA splicing

As we observed distinct m^6^A deposition inhibition by pre-mRNA splicing in individual flanking exons in the validation experiments (Figure 3), we further comprehensively investigated this exon heterogeneity of m^6^A deposition inhibition at a genome-wide scale. Towards this goal, we calculated the m^6^A probability change (ΔProbability) for the RAC sites located in all last exons after the last intron deletion in the gene for each gene in this study. The first 200 nucleotides of last exons were binned into 40 interval (5 nucleotides per interval). In each interval, the RAC site with maximum probability change was selected, and its corresponding ΔProbability was calculated as the ΔValue for the interval. Then based on the ΔValue and using the k-means clustering method, we clustered all the last exons into two groups: Cluster1 (C1) and Cluster2 (C2) (Fig. 4a for mouse, and Supplementary Fig. 5a for human). C1 exons were those highly enriched with the signal increased m^6^A sites (Fig. 4a for mouse, and Supplementary Fig. 5a for human), indicating C1 exons exhibited strong m^6^A deposition inhibition by pre-mRNA splicing. We found that 10.8% RAC sites in C1 exons showed increased m^6^A deposition (Fig. 4b for mouse, and Supplementary Fig. 5b for human), which was 3.5 fold of that in C2 exons (Fig. 4c for mouse, and Supplementary Fig. 5c for human). Furthermore, these splicing-repressed m^6^A sites (ΔProbability > 0.1) were enriched in the ~100 nt region of the C1 exons start (Fig. 4b for mouse, and Supplementary Fig. 5b for human). To further investigate these two distinct exon groups, we plotted their m^6^A levels before and after last intron deletion respectively. The m^6^A level at C1 exons was only mildly higher than that in C2 exons before last intron deletion in the gene (Fig. 4d,e,f for mouse, and Supplementary Fig. 5d,e,f for human). However, after last intron deletion in the gene, the m^6^A density increased sharply at C1 exons (about 5 fold), but not at C2 exons (Fig. 4e,f,g for mouse, and Supplementary Fig. 5e,f,g for human). To understand the underlying *cis*-element mechanism in the C1 and C2 exons, we compared the distribution of m^6^A enhancers and silencers around these splicing repressed m^6^A sites to that of RAC sites without m^6^A deposition change. The m^6^A enhancers were more enriched in the 50 nt downstream of the splicing repressed m^6^A sites in C1 exons (Fig. 4h,i for mouse, and Supplementary Fig. 5h,i for human), while the silencers were more avoided this region in comparison to these sites in C2 exons (Supplementary Fig. 7a,b for mouse, and Supplementary Fig. 7c,d for human). In addition, we found the RAC sites were strongly enriched (about 2 fold) in the ~100 nt region of exon start in C1 exons in comparison to that in C2 exons (Fig. 4j,k,l for mouse, and Supplementary Fig. 5j,k,l for human). Altogether, the m^6^A deposition inhibition by pre-mRNA splicing in last exons demonstrated a high heterogeneity: only a small proportion (mouse: 12.3%, 2339 out of 19045; human: 14.7%, 2681 out of 18209) of last exons exhibited strong inhibition, and these last exons contained a high density of RAC motif in the first 100 nt region of the last exon start.

**Fig. 4:**
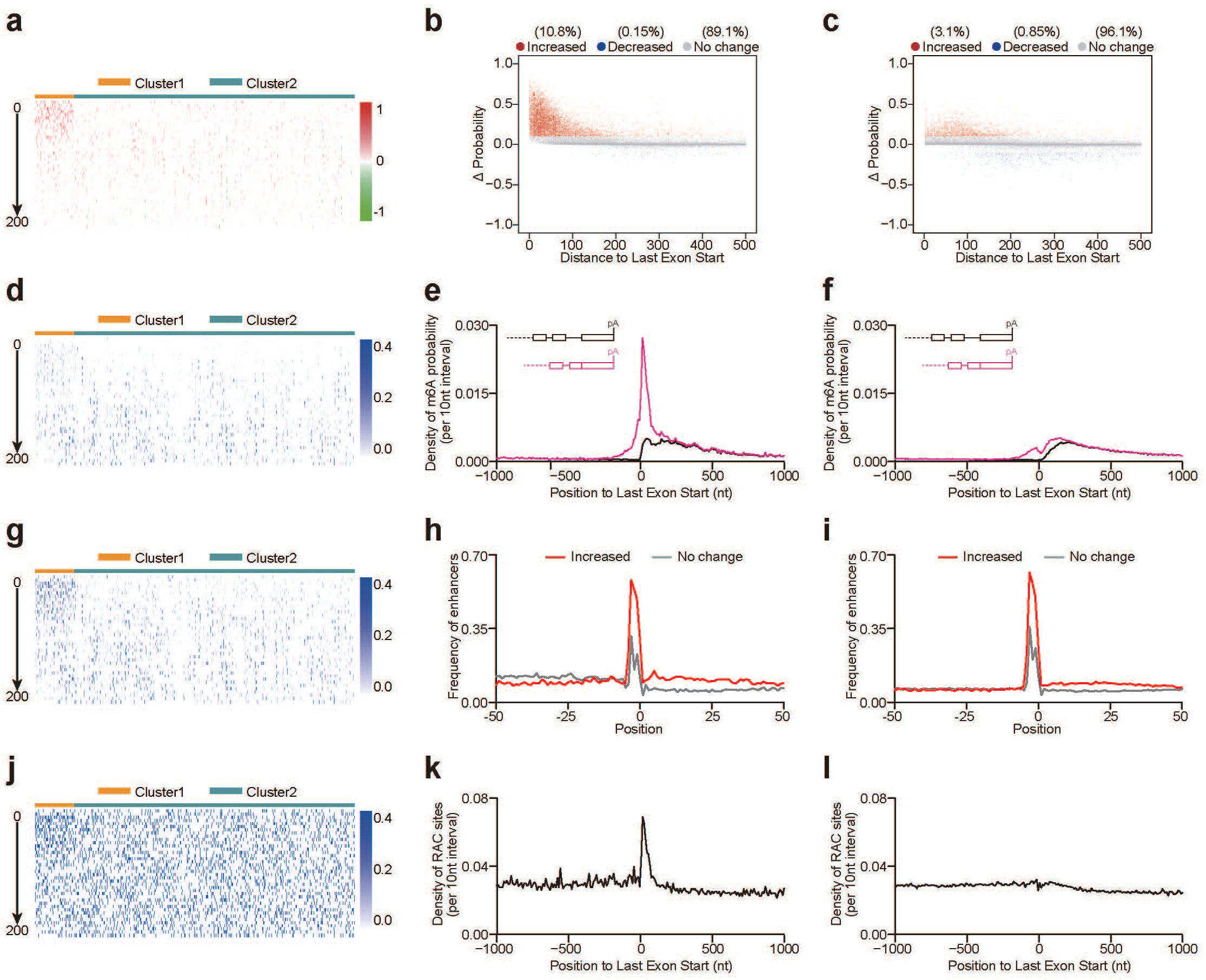
A proportion of last exons exhibit strong m^6^A deposition inhibition by pre-mRNA splicing. **a,d,g,j**, The heatmap visualized ΔProbability (Fig. 4a), m^6^A Probability (Fig. 4d), m^6^A Probability by last intron deletion (Fig. 4g), and counts of RAC sites (Fig. 4j) in the first 200 nt of last exon. The 200 nt was binned into 40 intervals (5 nt per interval). Genes were clustered (see details in Methods) into two clusters (Cluster1, Cluster2) based on ΔProbability. **b,c**, Positional plot of ΔProbability (Fig. 4b for Cluster1, Fig. 4c for Cluster2) for the RAC sites located in last exons. Red dots (Increased: n = 8700 for 2000 genes of Cluster1, n = 2509 for 2000 genes of Cluster2), blue dots (Decreased: n = 121 for 2000 genes of Cluster1, n = 691 for 2000 genes of Cluster2), and grey dots (no change: n = 72009 for 2000 genes of Cluster1, n = 78094 for 2000 genes of Cluster2) were those sites that had increased probability (> 0.1), decreased probability (< −0.1), or not change probability (|ΔProbability| <= 0.1) respectively by last intron deletion. **e,f**, The m^6^A density around last exon start (Fig. 4e for Cluster1, Fig. 4f for Cluster2) were compared for transcripts with full length (black line), last intron deletion (pink line). The value was calculated as the total probability value in a 10-nt interval divided by the total number of mRNAs in the interval. **h,i**, Positional plot for the frequency of top 50 enhancers (Fig. 4h for Cluster1, Fig. 4i for Cluster2) in mRNA sequences around the RAC sites. The sites were located in last exons, and the plots were compared between the increase sites (red line, ΔProbability > 0.1) and no change sites (grey line, |ΔProbability| <= 0.1). **k,l**, The density of RAC sites around last exon start (Fig. 4k for Cluster1, Fig. 4l for Cluster2). The value was calculated as the total number of RAC sites in a 10-nt interval divided by the total number of mRNAs in the interval.

### A small proportion of internal exons exhibit strong m^6^A deposition inhibition by pre-mRNA splicing

As our previous work demonstrated that last exons and internal exons might follow the same rule governing m^6^A deposition^23^, we speculated that internal exons might also demonstrate a high heterogeneity for m^6^A deposition inhibition by pre-mRNA splicing. Accordingly, for the RAC sites located in internal exons, we calculated using iM6A the m^6^A probability change (ΔProbability) after all introns were deleted in the gene, and applied the k-means method to cluster the internal exons into two groups: Cluster1 (C1) and Cluster2 (C2) (Fig. 5a for mouse, and Supplementary Fig. 6a for human). C1 exons were highly enriched with the increased m^6^A deposition sites (Fig. 5a for mouse, and Supplementary Fig. 6a for human), exhibiting strong m^6^A deposition inhibition by pre-mRNA splicing. In total, 69% of RAC sites in C1 exons showed increased m^6^A deposition (Fig. 5b for mouse, and Supplementary Fig. 6b for human), which was about 3 fold of that in C2 exons (Fig. 5c for mouse, and Supplementary Fig. 6c for human). Furthermore, the splicing-repressed m^6^A sites (ΔProbability > 0.1) were enriched in the ~100 nt region of C1 exon start (Fig. 5b, and Supplementary Fig. 6b). Before intron deletion in the gene, the m^6^A levels at internal exons were very low in both C1 and C2 exons (Fig. 5d,e,f for mouse, and Supplementary Fig. 6d,e,f for human). After intron deletion, the m^6^A density increased sharply at C1 exons, not at C2 exons (Fig. 5e,f,g for mouse, and Supplementary Fig. 6e,f,g for human). Consistent with the m^6^A enhancer and silencer distribution flanking RAC sites in last exons, the m^6^A enhancers were more enriched in the 50 nt downstream of increased sites in C1 exons (Fig. 5h,i for mouse, and Supplementary Fig. 6h,i for human), while the silencers tended to be avoided this region (Supplementary Fig. 7e,f for mouse, and Supplementary Fig. 7g,h for human). Lastly, the RAC sites were about 2 fold enriched in the ~100 nt region of exon start in C1 exons comparing to that in C2 exons (Fig. 5j,k,l for mouse, and Supplementary Fig. 6j,k,l for human). In summary, the m^6^A deposition inhibition by pre-mRNA splicing in internal exons also had a high heterogeneity, and a small proportion (mouse: 15%, 23473 out of 156514; human: 14.3%, 22805 out of 159561) of internal exons exhibited strong inhibition.

**Fig. 5:**
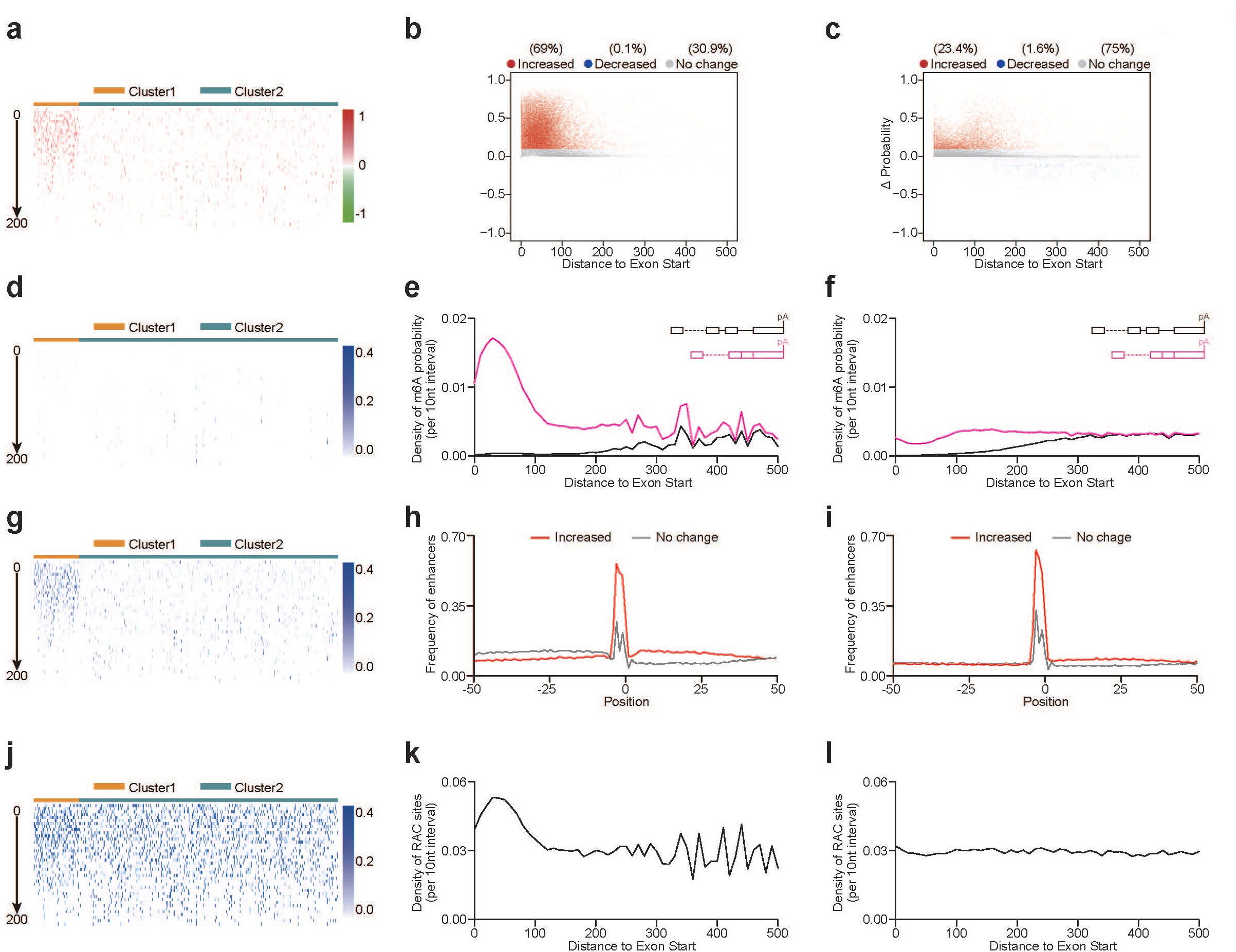
A proportion of internal exons exhibit strong m^6^A deposition inhibition by pre-mRNA splicing. **a,d,g,j**, The heatmap visualized ΔProbability (Fig. 5a), m^6^A Probability (Fig. 5d), m^6^A Probability by introns deletion (Fig. 5g), and counts of RAC sites (Fig. 5j) in the first 200 nt of internal exon. The 200 nt was binned into 40 intervals (5 nt per interval). Exons were clustered (see details in Methods) into two clusters (Cluster1, Cluster2) based on ΔProbability. **b,c**, Positional plot of ΔProbability (Fig. 5b for Cluster1, Fig. 5c for Cluster2) for the RAC sites located in internal exons. Red dots (Increased: n = 19368 for 5000 exons of Cluster1, n = 5458 for 5000 exons of Cluster2), blue dots (Decreased: n = 23 for 5000 exons of Cluster1, n = 363 for 5000 exons of Cluster2), and grey dots (no change: n = 8688 for 5000 exons of Cluster1, n = 17465 for 5000 exons of Cluster2) were those sites that had increased probability (> 0.1), decreased probability (< −0.1), or not change probability (|ΔProbability| <= 0.1) respectively by introns deletion. **e,f**, The m^6^A density at internal exon start (Fig. 5e for Cluster1, Fig. 5f for Cluster2) were compared for transcripts with full length (black line), introns deletion (pink line). The value was calculated as the total probability value in a 10-nt interval divided by the total number of mRNAs in the interval. **h,i**, Positional plot for the frequency of top 50 enhancers (Fig. 5h for Cluster1, Fig. 5i for Cluster2) in mRNA sequences around the RAC sites. The sites were in internal exons, and the plots were compared between the increase sites (red line, ΔProbability > 0.1) and no change sites (grey line, |ΔProbability| <= 0.1). **k,l**, The density of RAC sites at internal exon start (Fig. 5k for Cluster1, Fig. 5l for Cluster2). The value was calculated as the total number of RAC sites in a 10-nt interval divided by the total number of mRNAs in the interval.

### The m^6^A deposition inhibition by pre-mRNA splicing allows longer mRNA half-life

Since the pre-mRNA splicing inhibits m^6^A deposition at the nearby exons, one would expect an anti-correlation between the m^6^A deposition efficiency and the pre-mRNA splicing events (i.e. exon number) in the host genes. Indeed, in our minigene validation (Figure 3), we experimentally confirmed this hypothesis. To extend this finding at a genome-wide scale, we performed the scatter density plot between m^6^A/RAC ratio and the exon number in individual mRNAs, and observed a strongly negative correlation between the pre-mRNA splice events and m^6^A/RAC ratio (i.e. m^6^A deposition inhibition by pre-mRNA splicing) (Fig. 6a). Individual mRNAs with higher exon number had lower m^6^A deposition efficiency (Fig. 6a, and Supplementary Fig. 8a,b). Since a major function of m^6^A mRNA modification is to promote mRNA decay^9–12^, mRNAs with short half-lives (*T*_½_s < 5 h) had higher rate of m^6^A deposition, while mRNAs with longer half-lives (*T*_½_s of 5-10 h or >10 h) had a progressively lower rate of m^6^A deposition (Fig. 6b). However, this negative correlation between *T*_½_s and rate of m^6^A deposition vanished in mRNAs of *Mettl3* knockout mESCs (Fig. 6c), highlighting that this correlation is dependent on m^6^A. Similarly, mRNAs with short half-lives (*T*_½_s < 5 h) had fewer exons, while mRNAs with *T*_½_s of 5-10 h or >10 h had a progressively increased exon number (Fig. 6d, and Supplementary Fig. 8c). In addition, this correlation between *T*_½_s and exon numbers in individual mRNAs was also lost in *Mettl3* knockout mESCs (Fig. 6e, and Supplementary Fig. 8d).

**Fig. 6:**
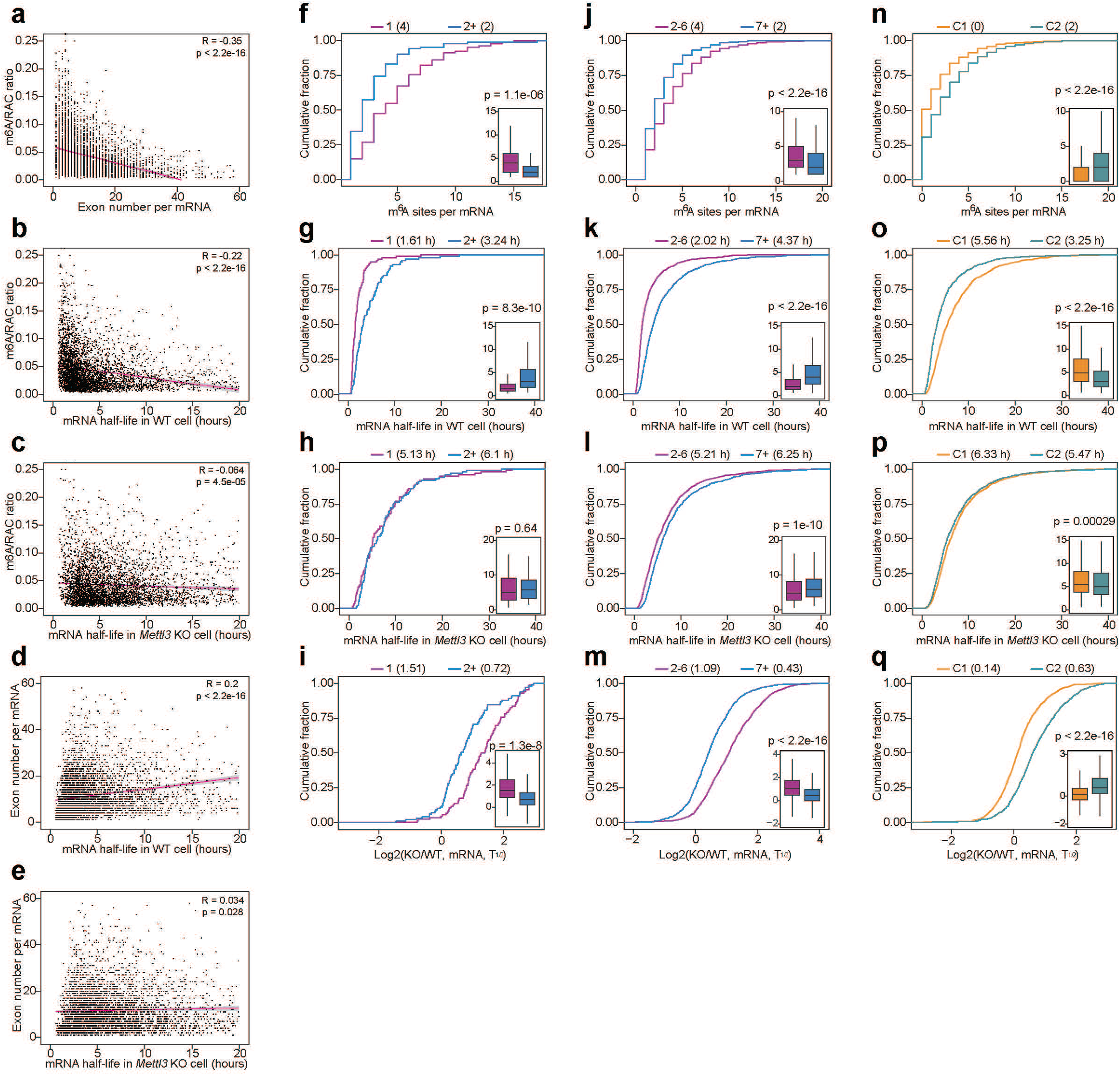
The m^6^A deposition inhibition by pre-mRNA splicing allows longer mRNA half-lives. **a**, The scatter density plot of m^6^A/RAC ratio for mRNA exon numbers for individual mRNAs (dots). **b-c**, The scatter density plot of m^6^A/RAC ratio for mRNA *T*_½_s (Fig. 6b for *Mettl3* WT mouse ES cells, Fig. 6c for *Mettl3* knockout mouse ES cells) for individual mRNAs (dots). **d-e**, The scatter density plot of exon numbers for mRNA *T*_½_s (Fig. 6b for *Mettl3* WT cells, Fig. 6c for *Mettl3* knockout cells) for individual mRNAs (dots). **f,g,h,i**, Cumulative distribution and boxplots (inset) showing m^6^A sites number (Fig. 6f), mRNA *T*_½_s in *Mettl3* WT cells (Fig. 6g), mRNA *T*_½_s in *Mettl3* knockout cells (Fig. 6h), and mRNA *T*_½_s changes upon global m^6^A loss (Fig. 6i) in single and multiple exon genes. Median and interquartile ranges were presented for the box plot. The p-values were calculated by Wilcoxon test. **j,k,l,m**, Cumulative distribution and boxplots (inset) showing m^6^A sites number (Fig. 6j), mRNA*T*_½_s in *Mettl3* WT cells (Fig. 6k), mRNA *T*_½_s in *Mettl3* knockout cells (Fig. 6l), and mRNA *T*_½_s changes upon global m^6^A loss (Fig. 6m) in genes with 2-6 exons and genes with >6 exons. Median and interquartile ranges were presented for the box plot. The p-values were calculated by Wilcoxon test. **n,o,p,q**, Cumulative distribution and boxplots (inset) showing m^6^A sites number (Fig. 6n), mRNA *T*_½_s in *Mettl3* WT cells (Fig. 6o), mRNA *T*_½_s in *Mettl3* knockout cells (Fig. 6p), and mRNA *T*_½_s changes upon global m^6^A loss (Fig. 6q) in genes of Cluster1 and genes of Cluster2. Median and interquartile ranges were presented for the box plot. The p-values were calculated by Wilcoxon test.

Having shown that m^6^A deposition efficiency is anti-correlated with pre-mRNA splicing events, it would be reasonable that mRNAs with fewer exons may have higher m^6^A levels. To verify this hypothesis, we compared the m^6^A level between single-exon and multiple exon genes by matching RAC sites in mRNAs (Fig. 6) or match cDNA length (Supplementary Fig. 8). We found that single-exon genes had higher number of m^6^A sites than multiple-exon genes (Fig. 6f, and Supplementary Fig. 8e). Since m^6^A negatively regulates mRNA half-life, these single-exon genes had shorter *T*_½_s (Fig. 6g, and Supplementary Fig. 8f) and greater *T*_½_s changes between *Mettl3* KO vs WT mESC cells (Fig. 6i, and Supplementary Fig. 8h). Moreover, the difference of *T*_½_s between single-exon and multiple-exon genes was lost upon global loss of m^6^A in *Mettl3* KO mESC cells (Fig. 6h, and Supplementary Fig. 8g). We performed a further analysis and found that mRNAs with 2-6 exons also had higher number of m^6^A sites than mRNAs with >=7 exons (Fig. 6j, and Supplementary Fig. 8i), and mRNAs with 2-6 exons also had shorter *T*_½_s (Fig. 6k, and Supplementary Fig. 8j) and greater *T*_½_s changes between Mettl3 KO vs WT mESC cells (Fig. 6m, and Supplementary Fig. 8l). Although *T*_½_s of mRNAs with 2-6 exons were shorter in *Mettl3* knockout mESCs (Fig. 6l, and Supplementary Fig. 8k), the difference of *T*_½_s (2-6 exons vs >=7 exons) was much smaller than that in *Mettl3* WT mESCs.

Since we discovered that m^6^A deposition was strongly inhibited in a small proportion of exons (C1 exons), we speculated that mRNAs with C1 exons would have lower m^6^A levels than these without C1 exons. As expected, mRNAs with C1 exons had fewer number of m^6^A sites (Fig. 6n, and Supplementary Fig. 8m), longer *T*_½_s (Fig. 6o, and Supplementary Fig. 8n) and smaller *T*_½_s changes between Mettl3 KO vs WT mESC cells (Fig. 6q, and Supplementary Fig. 8p). In addition, the difference of *T*_½_s (C1 vs C2) was almost lost upon global loss of m^6^A in Mettl3 KO mES cells (Fig. 6p, and Supplementary Fig. 8o). These data collectively demonstrate that pre-mRNA splicing inhibit<u>s</u> m^6^A deposition, allowing longer mRNA half-life.

### The m^6^A deposition inhibition by pre-mRNA splicing allows flexible protein coding

We had shown that RAC sites were enriched in the ~100 nt region of exon start in C1 exons. An open hypothesis is whether a distinct amino acid or codon usage exists in these exons. To test this hypothesis, we counted the codon usage for the first 30 codons (30 × 3 nt = 90 nt) in each exon, and also calculated its corresponding amino acid usage. We found that amino acids D, N, and T were the 3 mostly enriched in last exon of C1, while amino acids of S, P, and A were the 3 mostly avoided (Fig. 7a). Consistent with amino acids usage in last exon, D, N, and T were also enriched in internal exons of C1, while S, P, and A were avoided (Fig. 7b). The strong correlation of odds ratio (C1 vs C2) of amino acids usage (Fig. 7c) supported that last exons and internal exons follow the same amino acid usage bias to effect their m^6^A deposition^23^. As expected, the codons for D, N, T were enriched in C1 internal exons, while codons coding A, S, P were avoided (Fig. 7d,e). Moreover, the odds ratio (C1 vs C2) of codon usage also had strong correlation between last exon and internal exon (Fig. 7f). We noticed that sets of synonymous codons encoding the same amino acids had quite different codon usages in C1 versus C2 exons. For example, the GAC codon was more frequently used than synonymous codon GAT in C1 exons (Fig. 7g), and AAC codon was also more enriched than synonymous AAT codon (Fig. 7h). These data suggest that the m^6^A deposition inhibition by pre-mRNA splicing might allow flexible protein coding that could be needed in the C1 exons. Though these exons contained the biased amino acid and codon usage for specific protein coding and beyond, they didn’t appear to have the enriched m^6^A signal due to the m^6^A deposition inhibition by pre-mRNA splicing.

**Fig. 7:**
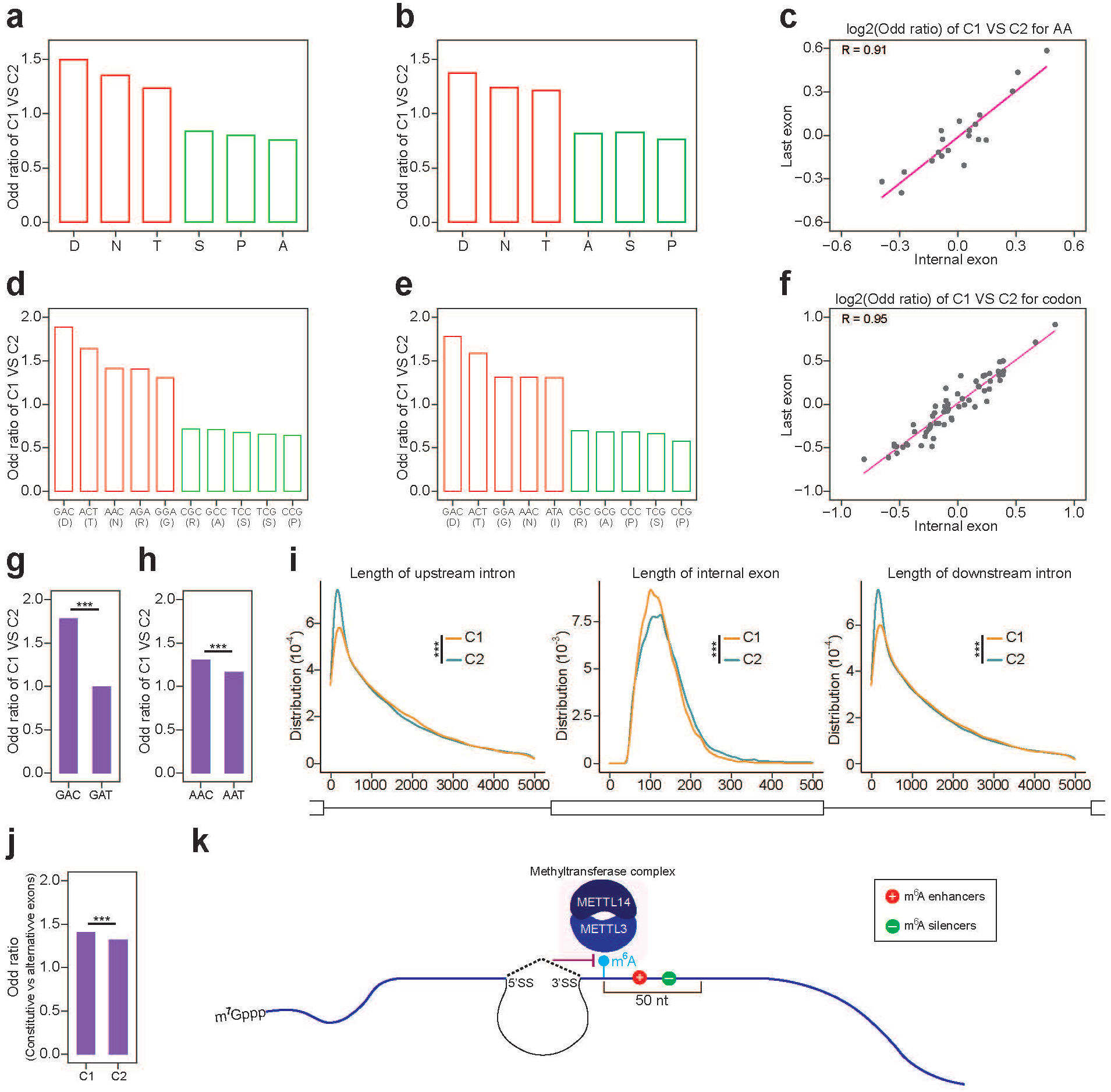
The m^6^A deposition inhibition by pre-mRNA splicing allows flexible protein coding. **a,b**, Bar plot of odd ratio for amino acids of Cluster1 vs Cluster2 in last exons (Fig. 7a) and internal exons (Fig. 7b). **c**, Scatter plot of the correlation for log2(odd ratio) of Cluster1 vs Clusters for amino acids in last exons and internal exons. Each gray dot was an amino acid. **d,e**, Bar plot of odd ratio for codons of Cluster1 vs Cluster2 in last exons (Fig. 7d) and internal exons (Fig. 7e). Its corresponding amino acids were also labeled. **f**, Scatter plot of the correlation for log2(odd ratio) of Cluster1 vs Clusters for codons in last exons and internal exons. Each gray dot was a codon. **g**, Bar plot of odd ratio for codons (GAC, GAT) encoding D amino acid of Cluster1 vs Cluster2 in internal exons. The p-values were calculated by Fisher-exact test (Significance: *** p < 0.001). **h**, Bar plot of odd ratio for codons (AAC, AAT) encoding N amino acid of Cluster1 vs Cluster2 in internal exons. The p-values were calculated by Fisher-exact test (Significance: *** p < 0.001). **i**, Density plot of internal exon length (middle panel), upstream intron length (left panel), and downstream intron length (right panel). The density was compared between Cluster1 and Cluster2. The p-values were calculated by the Kolmogorov–Smirnov test (Significance: *** p < 0.001). **j**, Bar plot of odd ratio for constitutive vs alternative exons in Cluster1 and Cluster2. The p-values were calculated by Fisher-exact test (Significance: *** p < 0.001). **k**, The site-specificity of m^6^A modification is determined by both local *cis*-elements within 50 nt downstream sequence and the intron inhibition to m^6^A deposition at nearby exons.

Besides the protein coding bias, we found that the length of C1 internal exons was shorter than C2 internal exons, while the length of its nearby introns including upstream and downstream intron was longer (Fig. 7i). In addition, C1 exons were more likely to be constitutive exons than alternative exons (Fig. 7j).

In summary, by *in silico* high-throughput mutational modeling and experimental validations, we found that pre-mRNA splicing inhibited the m^6^A deposition at nearby exons. The site-specificity of m^6^A deposition were influenced by both local cis-regulatory elements and this pre-mRNA splicing inhibition mechanism. Our work provides new insights into the mechanism of m^6^A site-specific deposition and its global distributional bias (Fig. 7k).

## Discussion

In this study, we explored the larger scale *cis*-regulatory mechanisms for m^6^A site specificity beyond the local *cis*-regulatory elements. By iM6A deep learning modeling, we uncovered that pre-mRNA splicing inhibited a proportion of m^6^A deposition at nearby exons. These findings were supported by experimental validations. Further, we revealed that the m^6^A deposition inhibition by pre-mRNA splicing exhibited a high degree of heterogeneity in different exons at genomic level, with a strong inhibition in a small group of exons. This m^6^A deposition inhibition by pre-mRNA splicing allows mRNA to have longer half-life. In addition, though some exons have biased amino acid and synonymous codon usage for their specific need for protein coding or beyond, these exons don’t appear to have higher m^6^A level due to this m^6^A deposition inhibition by pre-mRNA splicing.

Our findings that pre-mRNA splicing inhibited m^6^A deposition at the nearby exonic region close to splice sites and that the splicing-repressed m^6^A sites were enriched within the ~100 nt exonic region from either splice site of an exon could help us understand the regional bias for m^6^A modification in mRNAs. Given that most internal exons in vertebrate are short (average size < 150 nt)^30^, their exonic regions are mostly within the ~100 nt distance to a splice site and hence the m^6^A deposition is inhibited by pre-mRNA splicing in short internal exons. It could explain why m^6^As are relatively enriched in last exons, as well as long internal exons^20^. As last exon is composed of some coding region and most of the 3’UTR contains >70% of all m^6^A modification in mRNAs^20^, the pre-mRNA splicing inhibition on m^6^A deposition could focus the concentration of m^6^A signal on last exons and enable the complex and novel 3’UTR regulations involving m^6^A related RNA biology.

Vertebrate genes primarily consist of short exons separated by large introns while lower eukaryotes genes (yeast as an example) are made up of a large number of intronless genes or genes with long exons separated by small introns^31^. In yeast, m^6^A methylation occurs only during meiosis as the METTL3 yeast homolog IME4 expression is only expressed in this time period^33–35^. In mammals, the m^6^A deposition inhibition by pre-mRNA splicing may allow transcripts to have low methylation level in general despite the widespread expression of METTL3 across different tissues and cell types. In this study, we showed that C1 internal exons exhibit strong m^6^A deposition inhibition by pre-mRNA splicing. Comparing to other exons, these C1 exons tend to be shorter in length while being flanked longer 5’ and 3’ introns (Fig. 7i), suggesting the exon definition model could play an important role for these C1 exons. Furthermore, the finding that C1 internal exons tend to be constitutive exons not alternative exons (Fig. 7j), suggesting that the robust pre-mRNA splicing efficiency of constitutive exon may contribute to the pre-mRNA splicing inhibition of m^6^A methylation.

A major function of m^6^A is to promote mRNA decay^9–12^. We demonstrated that the m^6^A deposition efficiency has a strong anti-correlation with pre-mRNA splicing events, and mRNAs with higher exon number have lower m^6^A deposition efficiency. Thus, m^6^A deposition inhibition by pre-mRNA splicing enables transcripts with multiple exons to have long mRNA half-life. As this study has shown, in comparison to transcripts with multiple exons, transcripts with single exon have higher m^6^A levels and possess shorter *T*_½_s. Similarly, transcripts with lower exon number have higher number of m^6^A sites, as well as shorter *T*_½_s. Many important regulatory genes are intronless, including many immediate early genes (e.g. c-*Fos* gene) and important transcriptional factors (e.g. *Sox2* gene). The mRNAs of these genes are generally short-lived and have many m^6^As. Being intronless with more methylated sites, this leads to shorter half-life and lower activity, often appropriate for their evolved function to be able to response acutely to rapid environmental perturbations.

It has been well established that pre-mRNA splicing could influence mRNA half-life through the <u>n</u>on-sense <u>m</u>ediated <u>d</u>ecay (NMD) pathway^36^, and our finding that pre-mRNA splicing inhibited m^6^A deposition to increase mRNA half-life provided a completely new avenue for the regulation of pre-mRNA splicing on mRNA stability.

## Methods

### Modeling m^6^A deposition in pre-mRNA by iM6A

We pulled singularity container (tensorflow-19.01-py2) from NVIDA official website to create the environment for iM6A^23^, extra packages including biopython (1.76), scikit-learn (0.20.3), keras(2.0.5) were installed into external path by pip. The gene annotation tables (vM7 for mouse, v19 for human) were downloaded from GENCODE (https://www.gencodegenes.org/), and the longest transcript was extracted for each gene. The nucleotide sequence of pre-mRNA served as input, and the probability of each nucleotide being a m^6^A site was calculated by iM6A (Fig. 1a). For intron deletion, the sequences of the corresponding introns were deleted from the gene, and the m^6^A density around last exon start was compared between full length transcripts and the intron deletion control. For the RAC sites in exonic regions, the delta changes of m^6^A probability value (ΔProbability) after intron deletion were calculated. Then, the sites were categorized into three groups (increased, decreased and no change) based on ΔProbability (cutoff = 0.1). Positional plot and scatter plot were used to characterize ΔProbability distribution in exons.

### Positional plot of pentamers in sequences flanking m^6^A sites

For the RAC sites in last exon and second-to-last exon, we calculated their m^6^A probability change (ΔProbability) for last intron deletion by iM6A. The sites were categorized into three groups (increase, decrease and no change) based on ΔProbability (cutoff = 0.1). We extracted the 55 nt upstream and downstream sequences flanking the RAC sites in mRNA, and the pentamers were enumerated from the 5’ end to the 3’ end of the sequence. The m^6^A enhancers and silencers were quantified by iM6A through saturation mutation data analysis^23^. For positional plot, we counted the numbers of top 50 enhancers and top 50 silencers at each position of sequence. Then, the frequency of the enhancers or silencers were calculated. The plots were compared between the increased sites and no change sites. Similar strategy was applied to the RAC sites in internal exons.

### Conservation analysis of RAC sites

For the RAC sites in last exon and second-to-last exon, we calculated their m^6^A probability change (ΔProbability) for last intron deletion by iM6A. The RAC sites were categorized into three groups (increased, decreased and no change) based on ΔProbability (cutoff = 0.1). Those sites in degeneration position of synonymous codons were selected, and box plot was used to compare the PhyloP score between increased and no change sites. The p-values were determined by Wilcoxon test. Similar strategy was applied to the RAC sites in internal exons.

### Point mutation for 5’ and 3’ splice sites of last intron in pre-mRNA

For multi-exon genes (>=3 exons), its sequences of last introns were truncated to 200 nucleotides by keeping 100 nucleotides of intron start and intron end. Next, the 5’ splice site (donor: GT dinucleotide), 3’ splice site (acceptor: AG dinucleotide) of mini-introns were mutated to CA, TC respectively. In addition, the cryptic splice sites were predicted by SpliceAI^37^ for the sequence of second-to-last exon, mutated truncation intron and last exon. All of cryptic splice sites (Probability > 0.1) were also mutated (donor: mutated to CA; acceptor: mutated to TC). Finally, we only kept the genes (n = 2370) which had no new cryptic sites after this 1^st^ round of cryptical splice site point mutation according to SpliceAI, and iM6A was used to model the m^6^A deposition.

### Construction of the minigene

The backbone of minigene was a common retroviral GFP vector, and puromycin was the selection marker for stable cell line. *Gne*, and *Lrp12* were used as the two model genes for experimental validation. For each mRNA, the second-to-last exon was truncated to 100 nt by keeping the 100 nt exonic sequence upstream of the exon end, last intron was truncated to 200 nt by keeping the 100 nt intronic sequences at each end of the last intron, and last exon was truncated to 240 nt by preserving the 240nt downstream of the exon start. The AcGFP1 was in-frame fused to the second-to-last exon. To avoid <u>n</u>on-sense <u>m</u>ediated <u>d</u>ecay (NMD) effect, both genes have stop codon in the last exon. The detailed sequence for the *Gne* and *Lrp12* constructs are in the Supplementary Data1

### mRNA decay assay

The stable cell lines constantly expressing the minigenes were subjected to four time points (0, 3, 6, and 9 h) of post actinomycin D treatment (final concentration of 1 μg/mL; Sigma, no. A9415) treatment in three biological replicates. Total RNA of each sample was extracted and quantified by qRT-PCR. The normalized mRNA levels at 0 h were set to 100%. The *T*_½_ was determined as *ln*(2)/*k*, where *k* is the decay rate constant. The mRNA levels at different time points were fitted to a first-order exponential decay curve to calculate the *k*.

### m^6^A quantification by SELECT method

The constructs of minigenes were transfected to HEK-293T, and total RNA was extracted after 48 h. The elongation and ligation-based qPCR amplification method SELECT^29^ was used to quantify the m^6^A modification. For each RAC site in mRNA, the Ct value of m6A sites was first normalized to two non-RAC sites at each construct to calculate the m6A signal level for each site; the fold change of intensity for each m6A site was calculated by comparing their normalized Ct value differences for each m6A site between intron-containing and intron-deletion constructs. Oligos are listed in Supplementary Data2.

### Clustering exons based on ΔProbability of m^6^A by intron deletion

For the RAC sites located in last exons (Fig. 4 for mouse, and Supplementary Fig.5 for human), we calculated the delta changes of m^6^A probability value (ΔProbability) by last intron deletion. The first 200 nt of last exon was binned into 40 intervals (5 nt per interval). In each interval, the site with maximum of probability change was selected, while its corresponding ΔProbability was kept as the ΔValue for the interval. Exons then were clustered into two clusters (Cluster1: abbreviated C1, Cluster2: abbreviated C2) by k-means method based on the ΔValue. The heatmap visualized ΔValue (Fig. 4a), average m^6^A Probability (Fig. 4d), average m^6^A Probability after last intron deletion (Fig. 4g), and average count of RAC sites (Fig. 4j) in each interval. The same strategy was applied to cluster the internal exons upon all introns deletion (Fig. 5 for mouse, and Supplementary Fig. 6 for human).

### Correlation analysis between m^6^A and exon numbers

For each transcript, the m^6^A sites (Probability > 0.05) were predicted by iM6A, and total number of RAC sites in exons were also counted. Scatter density plot was used to visualize the correlation between m^6^A/RAC ratio and exon numbers (Fig. 6a). The R-value was calculated by Pearson Correlation Coefficient, and p-value was determined by two-sided t-test. In addition, the transcripts were binned based on exon numbers per mRNA, and boxplot was used to show the m^6^A/RAC ratio or m^6^A density (number of m^6^A sites per 100 nt) in each bin (Supplementary Fig. 8a,b).

### Correlation analysis between m^6^A and mRNA half-life

The mRNA half-lives data were downloaded from Gene Expression Omnibus repository under accession no.GSE86336, Scatter density plot was used to visualize the correlation between m^6^A/RAC ratio and mRNA half-lives (*T*_½_) in *Mettl3* WT (Fig. 6b) or knockout mouse ES cells (Fig. 6c). Similarly, the correlation between exon numbers per mRNA and mRNA *T*_½_s in *Mettl3* WT (Fig. 6d) or knockout cells (Fig. 6e) was plotted. In addition, the transcripts were binned based on exon numbers per mRNA, and boxplot was used to show the mRNA *T*_½_s in *Mettl3* WT (Supplementary Fig. 8c) or knockout cells (Supplementary Fig. 8d) for each bin. The R-value was calculated by Pearson Correlation Coefficient, and p-value was determined by two-sided t-test.

### Analysis of mRNA half-lives

The mRNA half-lives was compared for single-exon vs multiple-exons genes (Fig. 6f,g,h,i), 2-6 exons vs > 6 exons genes (Fig. 6j,k,l,m), C1 vs C2 genes (Fig. 6n,o,p,q). We matched the exact RAC sites (Fig. 6) or mRNA length (Supplementary Fig. 8) for transcripts, cumulative distribution and boxplots were used to show m^6^A sites number, mRNA *T*_½_s in *Mettl3* wild-type (WT) cells, mRNA *T*_½_s in *Mettl3* knockout (KO) cells, and mRNA *T*_½_s changes upon global m^6^A loss. Median and interquartile ranges were presented for the box plot. The p-values were calculated by Wilcoxon test.

### Comparison of amino acids or codons for C1 vs C2 exons

For the amino acids or codons in last exons or internal exons, we counted the number for each amino acid or codon. Only the genes expressed in mESCs were used (GSE86336). The frequency of amino acid or codon in C1 or C2 exons was calculated, and odd ratio of C1 vs C2 was computed. Fisher-exact test was used to evaluate the significance. Scatter plot was used to visualize the correlation of odds ratio between last exon and internal exon. The R-value was calculated by Pearson Correlation Coefficient.

## Data availability

The mRNA half-lives data were downloaded from the Gene Expression Omnibus repository under accession no.GSE86336.

## Code availability

The source code of iM6A is available at GitHub (https://github.com/ke-laboratory/iM6A-Splicing).

## Acknowledgments

We thank Dennis Weiss and members of Ke Laboratory and Ying Laboratory for comments, suggestions, and thoughtful discussions. Ke Laboratory and this research is funded by NIH/NIGMS Maximizing Investigators’ Research Award (MIRA) R35 Award (R35 GM133711 to S.K.), American Cancer Society Pilot Award (ACS-2019-Pilot-Ke/IRG-16-191-33/ IRG-21-136-36-IRG to S.K.) and the Jackson Laboratory Cancer Center New Investigator award from the NIH/NCI Cancer Center Support Grant (2 P30 CA034196-34 to S.K.).

## Author contributions

S.K., Z.L., and Z.Y. conceived and designed the study. Z.L. conducted the experiments and performed the data analysis. Q.M., S.S., N.L., and H.W. contributed to the test of experimental validation. S.K., and Z.L., wrote the manuscript. S.K. supervised the research.

## Competing interests

The authors declare that they have no competing interests.

## Notes

### Competing Interest Statement

The authors have declared no competing interest.

