## Supplemental Material for "Pre-mRNA splicing inhibits m^6^A deposition, allowing longer mRNA half-life and flexible protein coding"

**Supplementary Fig. 1: Deep learning modeling reveals last intron deletion promotes m$^6$A deposition at last exon and second-to-last exon (human data).**

**a**, The m$^6$A density of transcripts around last exon start were compared between full length (black line) and last intron deletion (pink line). The value was calculated as the total probability value in a 10-nt interval divided by the total number of mRNAs in the interval.

**b-c**, Positional plot of ΔProbability (Supplementary Fig. 1b) and scatter plot of probability (Supplementary Fig. 1c) for the RAC sites located in last exons. Red dots (Increased: n = 7510 for 4000 genes), blue dots (Decreased: n = 1958 for 4000 genes), and grey dots (no change: n = 161845 for 4000 genes) were those sites that had increased probability (> 0.1), decreased probability (< -0.1), or not change probability (IΔProbabilityI <= 0.1) respectively by last intron deletion.

**d-e**, Positional plot for the frequency of top 50 enhancers (Supplementary Fig. 1d), silencers (Supplementary Fig. 1e) in mRNA sequences around the RAC sites. The sites were located in last exons, and the plots were compared between the increase sites (red line, ΔProbability > 0.1) and no change sites (grey line, IΔProbabilityI <= 0.1).

**f**, Box plot of PhyloP score of latent m$^6$A sites or no change sites in last exons (n = 19248 or 129639). Median and interquartile ranges were presented for the box plot. The p-values were calculated by Wilcoxon test (Significance: *** p < 0.001).

**g**, The m$^6$A density of transcripts around second-to-last exon start were compared between full length (black line) and last intron deletion (pink line).

**h-i**, Positional plot of ΔProbability (Supplementary Fig. 1h) and scatter plot of probability (Supplementary Fig. 1i) for the RAC sites located in second-to-last exons. Red dots (Increased: n = 17760 for 17313 genes), blue dots (Decreased: n = 642 for 17313 genes), and grey dots (no change: n = 73973 for 17313 genes) were those sites that had increased probability (> 0.1), decreased probability (< -0.1), or not change probability (IΔProbabilityI <= 0.1) respectively by last intron deletion.

**j-k**, Positional plot for the frequency of top 50 enhancers (Supplementary Fig. 1j), silencers (Supplementary Fig. 1k) in mRNA sequences around the RAC sites. The sites were located in second-to-last exons, and the plots were compared between the increased sites (red line, ΔProbability > 0.1) and no change sites (grey line, IΔProbabilityI

<= 0.1).

**I**, Box plot of PhyloP score of latent m$^6$A sites or no change sites in second-to-last exons (n = 15625 or 57279). Median and interquartile ranges were presented for the box plot. The p-values were calculated by Wilcoxon test (Significance: *** $p < 0.001$).

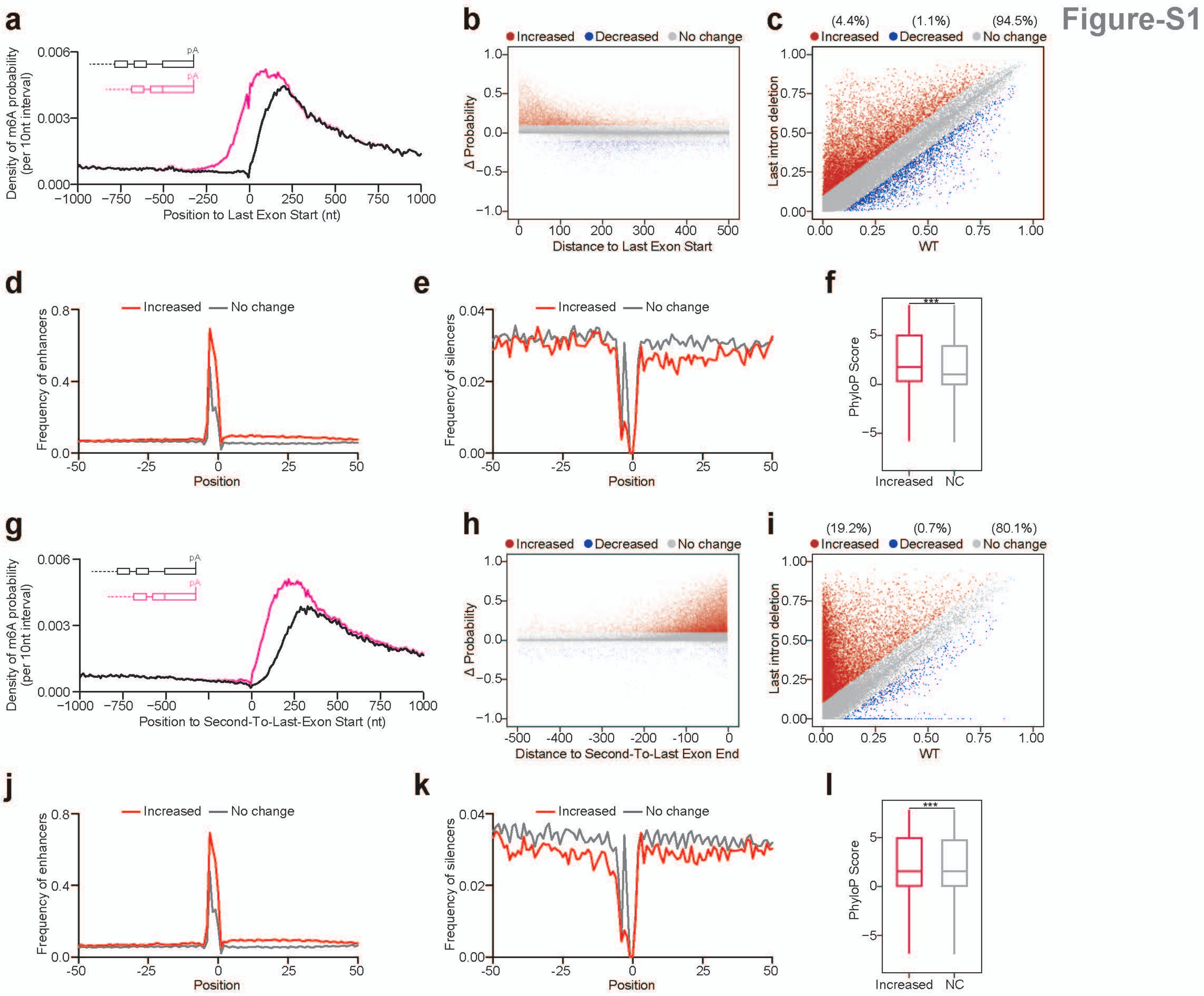
Figure-S1
a
Density of m6A probability (per 10nt interval)
Position to Last Exon Start (nt)
b
Increased Decreased No change
Δ Probability
Distance to Last Exon Start
c
(4.4%) (1.1%) (94.5%)
Increased Decreased No change
Last intron deletion
WT
d
Increased No change
Frequency of enhancers
Position
e
Increased No change
Frequency of silencers
Position
f
PhyloP Score
Increased NC
g
Density of m6A probability (per 10nt interval)
Position to Second-To-Last-Exon Start (nt)
h
Increased Decreased No change
Δ Probability
Distance to Second-To-Last Exon End
i
(19.2%) (0.7%) (80.1%)
Increased Decreased No change
Last intron deletion
WT
j
Increased No change
Frequency of enhancers
Position
k
Increased No change
Frequency of silencers
Position
l
PhyloP Score
Increased NC

**Supplementary Fig. 2: Deep learning modeling reveals introns deletion promotes $m^6A$ deposition at internal exons (human data).**

**a**, The $m^6A$ density of transcripts around last exon start were compared between full length (black line) and all introns deletion (pink line).

**b-d**, Positional plot of ΔProbability (Supplementary Fig. 2b,c) and scatter plot of probability (Supplementary Fig. 2d) for the RAC sites located in internal exons. Red dots (Increased: n = 20087 for 1500 genes), blue dots (Decreased: n = 1741 for 1500 genes), and grey dots (no change: n = 63192 for 1500 genes) were those sites that had increased probability (> 0.1), decreased probability (< -0.1), or not change probability (IΔProbabilityI <= 0.1) respectively by all introns deletion.

**e-f**, Positional plot for the frequency of top 50 enhancers (Supplementary Fig. 2e), silencers (Supplementary Fig. 2f) in mRNA sequences around the RAC sites. The sites were located in internal exons, and the plots were compared between the increased sites (red line, ΔProbability > 0.1) and no change sites (grey line, IΔProbabilityI <= 0.1).

**g**, Box plot of PhyloP score of latent $m^6A$ sites or no change sites in internal exons (n = 192830 or 605033). Median and interquartile ranges were presented for the box plot. The p-values were calculated by Wilcoxon test (Significance: *** $p < 0.001$).

**h-i**, The $m^6A$ density around last exon start were compared between with the full length transcripts (black line) and its intron truncation control (pink line). (Supplementary Fig. 2h for last intron truncation, Supplementary Fig. 2i for all introns truncation).

Figure-S2

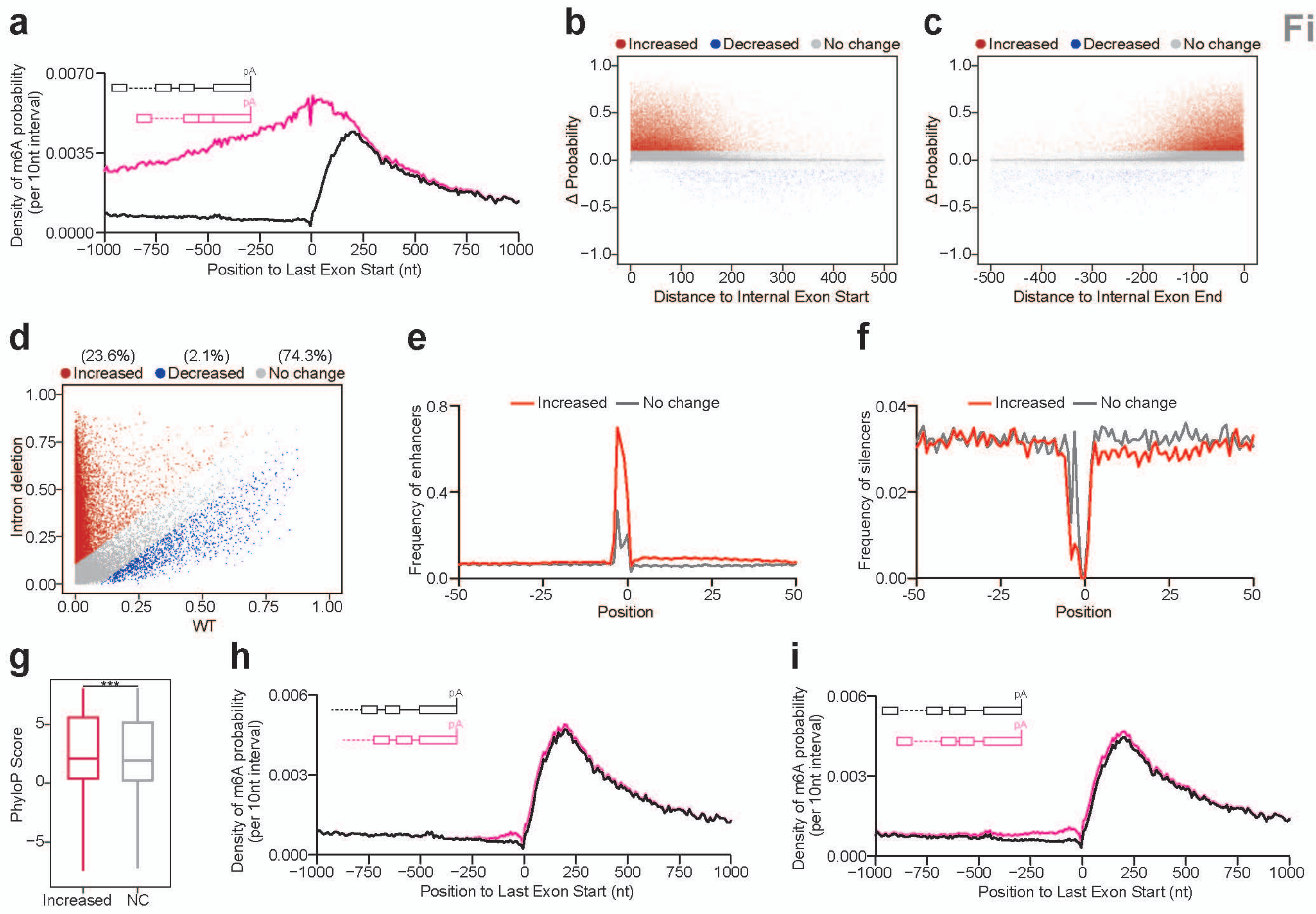
a
Density of m6A probability (per 10nt interval)
Position to Last Exon Start (nt)
b
Increased
Decreased
No change
Δ Probability
Distance to Internal Exon Start
c
Increased
Decreased
No change
Δ Probability
Distance to Internal Exon End
d
(23.6%) Increased
(2.1%) Decreased
(74.3%) No change
Intron deletion
WT
e
Increased
No change
Frequency of enhancers
Position
f
Increased
No change
Frequency of silencers
Position
g
***
PhyloP Score
Increased
NC
h
Density of m6A probability (per 10nt interval)
Position to Last Exon Start (nt)
i
Density of m6A probability (per 10nt interval)
Position to Last Exon Start (nt)

**Supplementary Fig. 3: Modeling $m^6A$ deposition in pre-mRNA with mini last intron**

**a**, The $m^6A$ density (10 nt interval) around last exon start were compared among the full length transcripts (black line) and their last intron truncated to 200 nt control (green line) and last intron deletion control (pink line).

**b-c**, The $m^6A$ density around last exon start (Supplementary Fig. 3b) or second-to-last exon end (Supplementary Fig. 3c) were compared among the full length transcripts (black line) and their last intron truncated to 200 nt control (green line) and last intron deletion control (pink line).

### Figure-S3

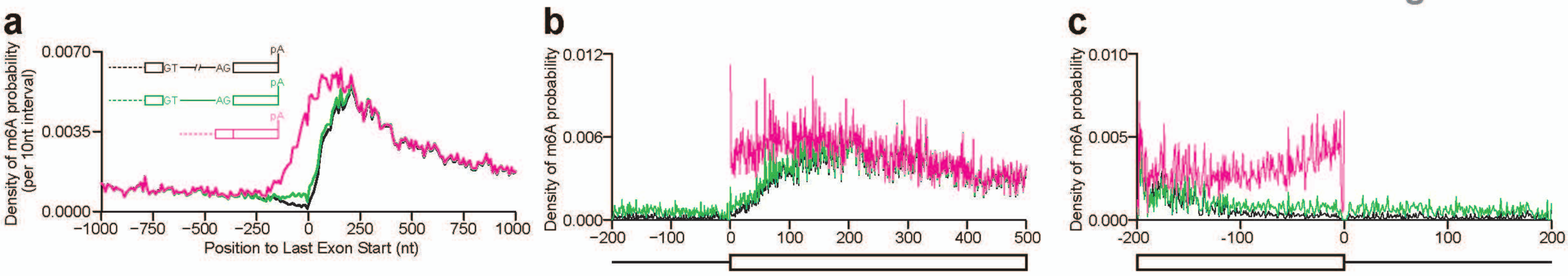
a
b
c
Density of m6A probability (per 10nt interval)
0.0070
0.0035
0.0000
GT
AG
pA
−1000
−750
−500
−250
0
250
500
750
1000
Position to Last Exon Start (nt)
Density of m6A probability
0.012
0.006
0.000
−200
−100
0
100
200
300
400
500
0.010
0.005
0.000
-200
-100
0
100
200

**Supplementary Fig. 4: Experimental validation of intron repression on m$^6$A deposition**

**a**, iM6A modeled methylation level (deposition probability) for each RAC site in minigene mRNA (left panel for *Lrp12*, and right panel for *Gne*). The m$^6$A level for constructs of intron-containing, and intron-deletion were marked by black and red colors. Each RAC site in mRNA was labeled as pink line.

**b**, The dot plot of the iM6A modeled ΔMethylation level for each RAC site in mRNA. ΔMethylation level was calculated for the m$^6$A signal difference of RAC sites between intron-containing and intron-deletion mRNAs. The mean value (0.197) was shown as the dotted pink line, and p-value was calculated by one-sample t-test for m$^6$A signal increase vs. no change.

**c-d**, The bar plot of qPCR ΔΔCt showing SELECT results for detecting the m$^6$A sites in mRNA. The constructs of minigenes were shown, and RAC sites in *Lrp12* (Supplementary Fig. 4c) or *Gne* (Supplementary Fig. 4d) were marked as pink line. Data were presented as mean ± SD, the p-values were calculated by Student t-test (Significance: *** $p < 0.001$, ** $p < 0.01$, * $p < 0.05$).

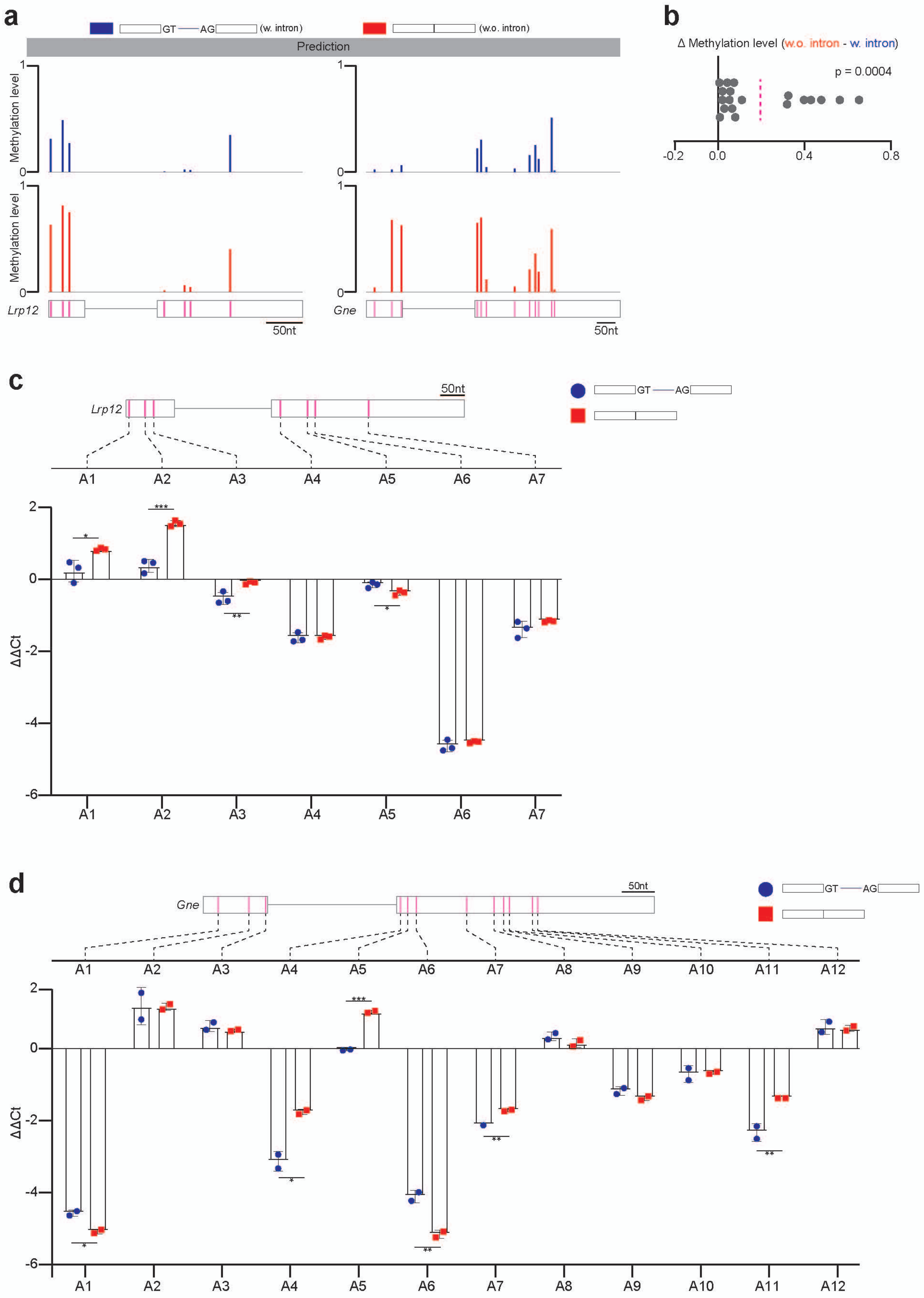
a
GT AG (w. intron)
(w.o. intron)
Prediction
Methylation level
1
0
Lrp12
Gne
50nt
b
Δ Methylation level (w.o. intron - w. intron)
p = 0.0004
-0.2
0.0
0.4
0.8
c
Lrp12
50nt
GT AG
A1
A2
A3
A4
A5
A6
A7
ΔΔCt
2
0
-2
-4
-6
d
Gne
50nt
A8
A9
A10
A11
A12

**Supplementary Fig. 5: A proportion of last exons exhibit strong m$^6$A deposition inhibition by pre-mRNA splicing (human data).**

**a,d,g,j**, The heatmap visualized ΔProbability (Supplementary Fig. 5a), m$^6$A Probability (Supplementary Fig. 5d), m$^6$A Probability by last intron deletion Supplementary Fig. 5g), and counts of RAC sites (Supplementary Fig. 5j) in the first 200 nt of last exon. The 200 nt was binned into 40 intervals (5 nt per interval). Genes were clustered (see details in Methods) into two clusters (Cluster1, Cluster2) based on ΔProbability.

**b,c**, Positional plot of ΔProbability (Supplementary Fig. 5b for Cluster1, Supplementary Fig. 5c for Cluster2) for the RAC sites located in last exon. Red dots (Increased: n = 10261 for 2000 genes of Cluster1, n = 2480 for 2000 genes of Cluster2), blue dots (Decreased: n = 242 for 2000 genes of Cluster1, n = 963 for 2000 genes of Cluster2), and grey dots (no change: n = 76388 for 2000 genes of Cluster1, n = 78404 for 2000 genes of Cluster2) were those sites that had increased probability (> 0.1), decreased probability (< -0.1), or not change probability (IΔProbabilityI <= 0.1) respectively by last intron deletion.

**e,f**, The m$^6$A density around last exon start (Supplementary Fig. 5e for Cluster1, Supplementary Fig. 5f for Cluster2) were compared between the full length transcripts (black line) and the last intron deletion control (pink line). The value was calculated as the total probability value in a 10-nt interval divided by the total number of mRNAs in the interval.

**h,i**, Positional plot for the frequency of top 50 enhancers (Supplementary Fig. 5h for Cluster1, Supplementary Fig. 5i for Cluster2) in mRNA sequences around the RAC sites. The sites were in last exons, and the plots were compared between the increased sites (red line, ΔProbability > 0.1) and no change sites (grey line, IΔProbabilityI <= 0.1).

**k,l**, The density of RAC sites around last exon start (Supplementary Fig. 5k for Cluster1, Supplementary Fig. 5l for Cluster2). The value was calculated as the total number of RAC sites in a 10-nt interval divided by the total number of mRNAs in the interval.

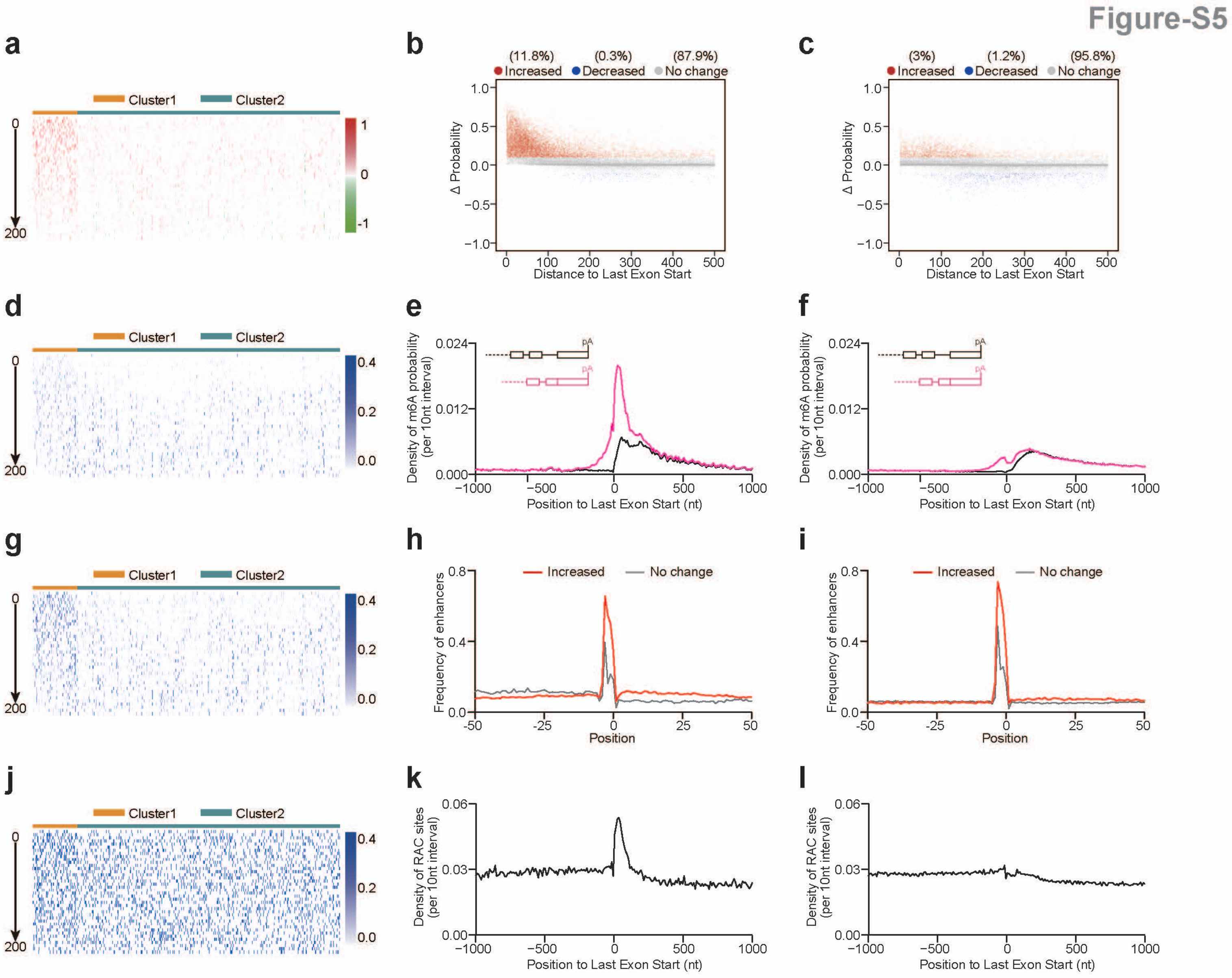
Figure-S5
a
Cluster1
Cluster2
0
200
1
0
-1
b
(11.8%)
Increased
(0.3%)
Decreased
(87.9%)
No change
Δ Probability
Distance to Last Exon Start
c
(3%)
Increased
(1.2%)
Decreased
(95.8%)
No change
Δ Probability
Distance to Last Exon Start
d
e
Density of m6A probability (per 10nt interval)
Position to Last Exon Start (nt)
pA
f
Density of m6A probability (per 10nt interval)
Position to Last Exon Start (nt)
g
h
Increased
No change
Frequency of enhancers
Position
i
Increased
No change
Frequency of enhancers
Position
j
k
Density of RAC sites (per 10nt interval)
Position to Last Exon Start (nt)
l
Density of RAC sites (per 10nt interval)
Position to Last Exon Start (nt)

**Supplementary Fig. 6: A proportion of internal exons exhibit strong m$^6$A deposition inhibition by pre-mRNA splicing (human data).**

**a,d,g,j**, The heatmap visualized ΔProbability (Supplementary Fig. 6a), m$^6$A Probability (Supplementary Fig. 6d), m$^6$A Probability by introns deletion (Supplementary Fig. 6g), and counts of RAC sites (Supplementary Fig. 6j) in the first 200 nt of internal exon. The 200 nt was binned into 40 intervals (5 nt per interval). Exons were clustered (see details in Methods) into two clusters (Cluster1, Cluster2) based on ΔProbability.

**b,c**, Positional plot of ΔProbability (Supplementary Fig. 6b for Cluster1, Supplementary Fig. 6c for Cluster2) for the RAC sites located in internal exons. Red dots (Increased: n = 19181 for 5000 exons of Cluster1, n = 4781 for 5000 exons of Cluster2), blue dots (Decreased: n = 55 for 5000 exons of Cluster1, n = 701 for 5000 exons of Cluster2), and grey dots (no change: n = 15235 for 5000 exons of Cluster1, n = 24133 for 5000 exons of Cluster2) were those sites that had increased probability (> 0.1), decreased probability (< -0.1), or not change probability (|ΔProbability| <= 0.1) respectively by introns deletion.

**e,f**, The m$^6$A density at internal exon start (Supplementary Fig. 6e for Cluster1, Supplementary Fig. 6f for Cluster2) were compared between the full length transcripts (black line) and the introns deletion control (pink line). The value was calculated as the total probability value in a 10-nt interval divided by the total number of mRNAs in the interval.

**h,i**, Positional plot for the frequency of top 50 enhancers (Supplementary Fig. 6h for Cluster1, Supplementary Fig. 6i for Cluster2) in mRNA sequences around the RAC sites. The sites were in internal exons, and the plots were compared between the increased sites (red line, ΔProbability > 0.1) and no change sites (grey line, |ΔProbability| <= 0.1).

**k,l**, The density of RAC sites at internal exon start (Supplementary Fig. 6k for Cluster1, Supplementary Fig. 6l for Cluster2). The value was calculated as the total number of RAC sites in a 10-nt interval divided by the total number of mRNAs in the interval.

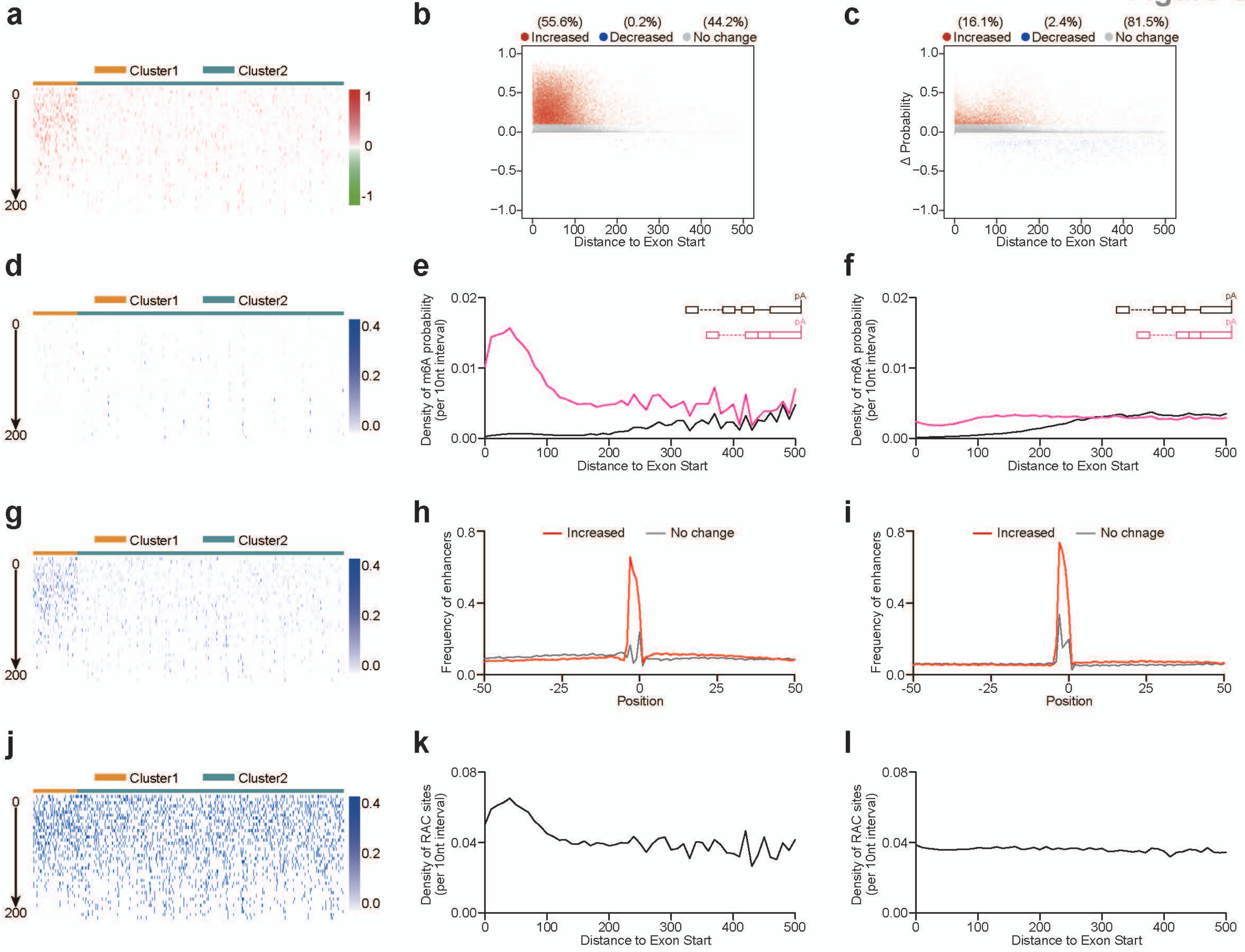
a
Cluster1
Cluster2
0
200
1
0
-1
b
(55.6%)
Increased
(0.2%)
Decreased
(44.2%)
No change
Distance to Exon Start
c
(16.1%)
Increased
(2.4%)
Decreased
(81.5%)
No change
Δ Probability
Distance to Exon Start
d
e
Density of m6A probability
(per 10nt interval)
Distance to Exon Start
pA
f
Density of m6A probability
(per 10nt interval)
Distance to Exon Start
g
h
Increased
No change
Frequency of enhancers
Position
i
Increased
No chnage
Frequency of enhancers
Position
j
k
Density of RAC sites
(per 10nt interval)
Distance to Exon Start
l
Density of RAC sites
(per 10nt interval)
Distance to Exon Start

**Supplementary Fig. 7: m$^6$A silencers avoids 50 nt downstream of latent sites.**

**a,b,c,d**, Positional plot for the frequency of top 50 silencers (Supplementary Fig. 7a,c for Cluster1 in mouse or human, Supplementary Fig. 7b,d for Cluster2 in mouse or human) in mRNA sequences around the RAC sites. These sites were in last exons, and the plots were compared between the increased sites (red line, ΔProbability > 0.1) and no change sites (grey line, IΔProbabilityI <= 0.1).

**e,f,g,h**, Positional plot for the frequency of top 50 silencers (Supplementary Fig. 7e,g for Cluster1 in mouse or human, Supplementary Fig. 7f,h for Cluster2 in mouse or human) in mRNA sequences around the RAC sites. These sites were located in internal exons, and the plots were compared between the increased sites (red line, ΔProbability > 0.1) and no change sites (grey line, IΔProbabilityI <= 0.1).

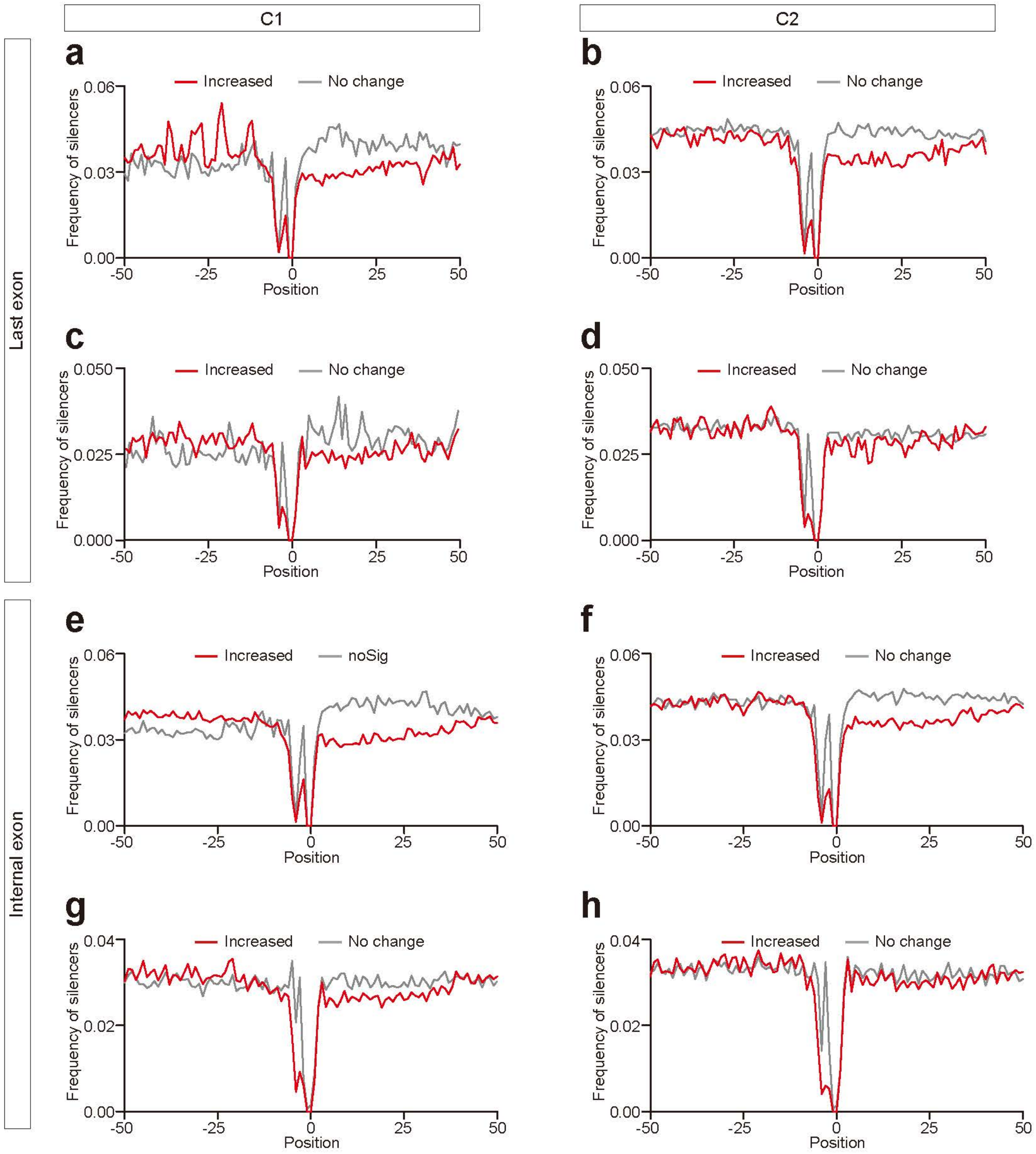
C1
C2
Last exon
Internal exon
a
b
c
d
e
f
g
h
Increased
No change
noSig
Frequency of silencers
Position
0.06
0.03
0.00
0.050
0.025
0.000
0.04
0.02
-50
-25
0
25
50

**Supplementary Fig. 8: The m$^6$A deposition inhibition by pre-mRNA splicing allows longer mRNA half-lives.**

**a,b,c,d**, Boxplots demonstrating the correlation between exon number per mRNA and m$^6$A/RAC ratio (Supplementary Fig. 8a), m$^6$A density per 100 nt (Supplementary Fig. 8b), mRNA $T_{1/2}$s in *Mettl3* WT mouse ES cells (Supplementary Fig. 8c), mRNA $T_{1/2}$s in *Mettl3* knockout mouse ES cells (Supplementary Fig. 8d). Median and interquartile ranges were presented for the box plot, and each dot represented a unique mRNA.

**e,f,g,h**, Cumulative distribution and boxplots (inset) showing m$^6$A sites number (Supplementary Fig. 8e), mRNA $T_{1/2}$s in *Mettl3* WT mouse ES cells (Supplementary Fig. 8f), mRNA $T_{1/2}$s in *Mettl3* knockout cells (Supplementary Fig. 8g), and mRNA $T_{1/2}$s changes upon global m$^6$A loss (Supplementary Fig. 8h) in single and multiple exon genes. Median and interquartile ranges were presented for the box plot. The p-values were calculated by Wilcoxon test.

**i,j,k,l**, Cumulative distribution and boxplots (inset) showing m$^6$A sites number (Supplementary Fig. 8i), mRNA $T_{1/2}$s in *Mettl3* WT mouse ES cells (Supplementary Fig. 8j), mRNA $T_{1/2}$s in *Mettl3* knockout cells (Supplementary Fig. 8k), and mRNA $T_{1/2}$s changes upon global m$^6$A loss (Supplementary Fig. 8l) in genes with 2-6 exons and genes with > 6 exons. Median and interquartile ranges were presented for the box plot. The p-values were calculated by Wilcoxon test.

**m,n,o,p**, Cumulative distribution and boxplots (inset) showing m$^6$A sites number (Supplementary Fig. 8m), mRNA $T_{1/2}$s in *Mettl3* WT mouse ES cells (Supplementary Fig. 8n), mRNA $T_{1/2}$s in *Mettl3* knockout cells (Supplementary Fig. 8o), and mRNA $T_{1/2}$s changes upon global m$^6$A loss (Supplementary Fig. 8p) in genes of Cluster1 and genes of Cluster2. Median and interquartile ranges were presented for the box plot. The p-values were calculated by Wilcoxon test.

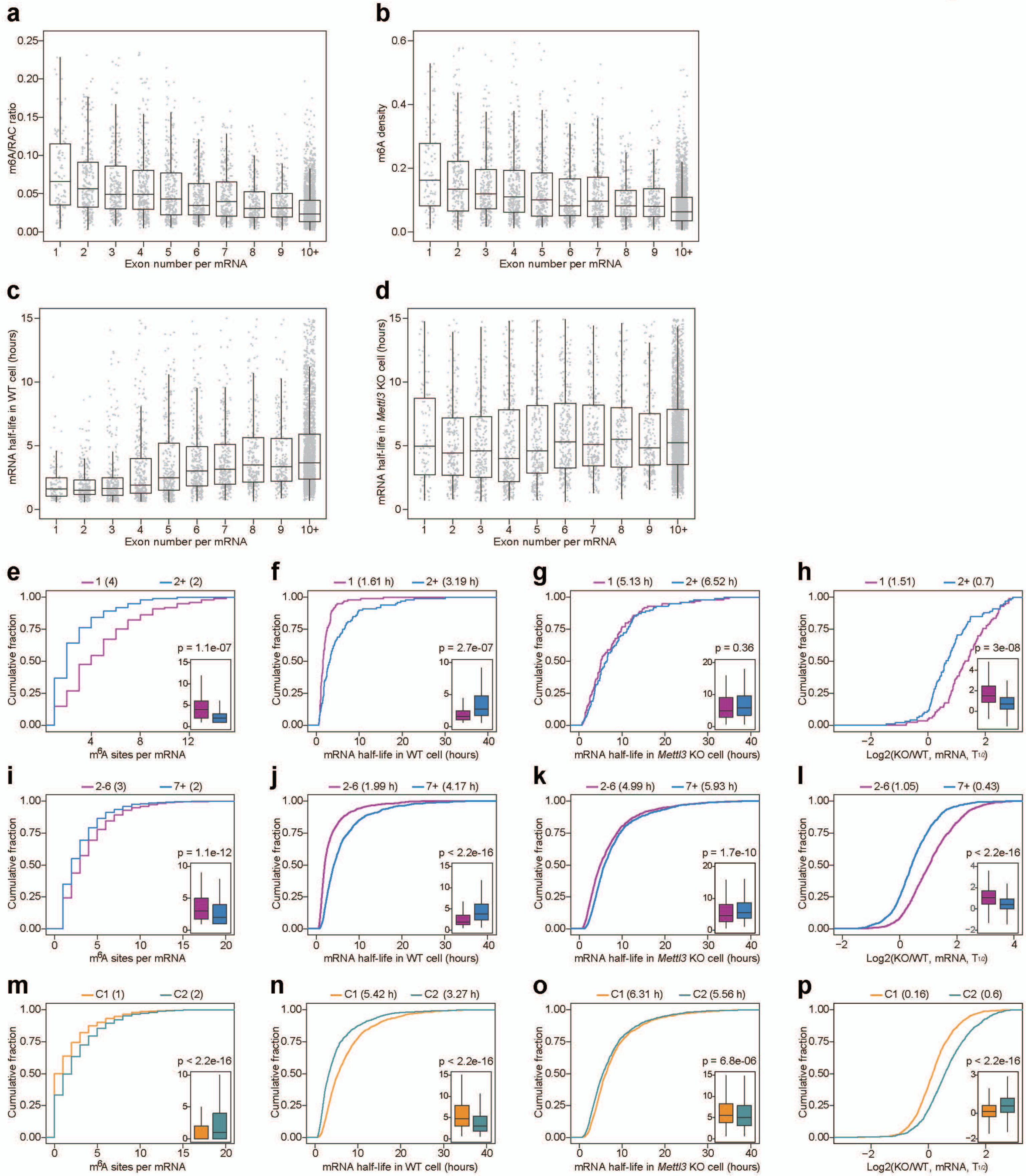

a
m6A/RAC ratio
Exon number per mRNA
b
m6A density
Exon number per mRNA
c
mRNA half-life in WT cell (hours)
Exon number per mRNA
d
mRNA half-life in Mettl3 KO cell (hours)
Exon number per mRNA
e
1 (4)
2+ (2)
p = 1.1e-07
Cumulative fraction
m6A sites per mRNA
f
1 (1.61 h)
2+ (3.19 h)
p = 2.7e-07
mRNA half-life in WT cell (hours)
g
1 (5.13 h)
2+ (6.52 h)
p = 0.36
mRNA half-life in Mettl3 KO cell (hours)
h
1 (1.51)
2+ (0.7)
p = 3e-08
Log2(KO/WT, mRNA, T1/2)
i
2-6 (3)
7+ (2)
p = 1.1e-12
m6A sites per mRNA
j
2-6 (1.99 h)
7+ (4.17 h)
p < 2.2e-16
mRNA half-life in WT cell (hours)
k
2-6 (4.99 h)
7+ (5.93 h)
p = 1.7e-10
mRNA half-life in Mettl3 KO cell (hours)
l
2-6 (1.05)
7+ (0.43)
p < 2.2e-16
Log2(KO/WT, mRNA, T1/2)
m
C1 (1)
C2 (2)
p < 2.2e-16
m6A sites per mRNA
n
C1 (5.42 h)
C2 (3.27 h)
p < 2.2e-16
mRNA half-life in WT cell (hours)
o
C1 (6.31 h)
C2 (5.56 h)
p = 6.8e-06
mRNA half-life in Mettl3 KO cell (hours)
p
C1 (0.16)
C2 (0.6)
p < 2.2e-16
Log2(KO/WT, mRNA, T1/2)
